# Measurements of Contractility in Actin Convergence Zone and KIF11-Inhibited Mitotic Spindles

**DOI:** 10.1101/2024.06.28.601275

**Authors:** Alexandre Matov

**Affiliations:** DataSet Analysis LLC, 155 Jackson St, San Francisco, CA 94111, United States

**Keywords:** Contractile Cytoskeletal Structures, Mitotic Spindle, Actin Meshworks, Growth Cones, Monastrol, S-trityl-L-cysteine, Blebbistatin, Tropomyosin Isoforms, Weak Actin Binders, Instantaneous Flow Tracking Algorithm

## Abstract

**Introduction:** The complex dynamics of cytoskeletal meshworks make them a difficult subject of study. With the advent of fluorescent speckle microscopy (FSM) and other technological advances in microscopy techniques, much more is now known about how the filamentous actin (F-actin) and microtubule (MT) networks work within cells to give rise to the vast array of functions which require them. A current challenge to the imaging field is to improve the utility and accuracy of the computational approaches required to analyze large and complex imaging datasets. Here, we present the results of a computational method that, when applied to FSM time-lapse series, can capture the instantaneous state of the rapidly changing, dense, and multi-directional speckle flows often exhibited by cytoskeletal dynamics in living systems. Re-analysis of previously published FSM image sets demonstrates that this method, which we call the Instantaneous Flow Tracking Algorithm (IFTA), can accurately detect speckle flows in mitotic spindles and F-actin meshworks, even in regions of high dynamicity of overlapping, anti-parallel flows where previous methods failed.

**Methods:** The anti-parallel flow at the metaphase plate of the mitotic spindle is a well-known phenomenon during the initial stages of chromosome segregation and it has been measured by several approaches, mostly in stationary spindles which do not exhibit any polar rotation. The mitotic spindle is the target of many cancer and neurodegenerative drugs and as such, there has been many attempts at inhibiting its basic functions with the objective of preventing chromosome segregation and the formation of new daughter cells. Such attempts have largely been focused on the inhibition of the action of MT plus-end directed motors, for instance the kinesin KIF11. Spindles with inhibited kinesins have been thought to exhibit no MT flux, however IFTA measured regional flux of up to 2.7 µm/min, which reveals the activity of potent secondary flux mechanisms. These results suggest novel, additional, approaches toward arresting cells in mitosis during patient treatment.

**Results:** The traditional tracking methods erroneously measure zero flux in areas where contractile F-actin flows meet, denoted as a “convergence zone” and commonly found in the lamella of motile cells and the neck of growth cones. When IFTA was used to analyze FSM datasets derived from these structures, we detected high rates of protein turnover, anti-parallel speckle motion, and fast flux of actin subunits in both directions in the same “convergence zones”. This demonstrates the presence of a molecular machinery based on contractility in the lamella/lamellipodium of migrating cells and at the base of growing neurons, which can be exploited in the clinic. When applied to FSM data of migrating kangaroo rat kidney epithelial PtK1 cells overexpressing different isoforms of the actin-based motor tropomyosin, IFTA revealed distinct, isoform-specific effects on contractile F-actin flows. Specifically, we found that decreased affinity between tropomyosin and F-actin correlated with an increase in speckle velocity heterogeneity. Such quantitative imaging analysis offers the ability to reliably test novel therapeutics *ex vivo*.

**Conclusions:** The main discoveries presented in the manuscript are related to the ability of the cell to adapt to a range of biochemical and mechanical perturbations. As cell therapy is becoming the forefront of precision medicine, it would be critical to anticipate the mechanisms of action of engineered immune cells. The pathways activated during therapy can be pinpointed by measuring its effects on the target proteins in patient-derived living cells. The clinical application of the approach outlined in this manuscript pertains to analyzing drug resistance in cancer therapy and the treatment of neurodegeneration. Our hypothesis is that targeting actin in resistant tumors could sensitize cancer cells to tubulin inhibitors. If this proves true, it will have implications in the clinic.

## INTRODUCTION

Quantitative analysis of fluorescent speckle microscopy (FSM) (Waterman-Storer et al., 1998) datasets provides a powerful means to probe molecular mechanisms, but resolving speckle motion poses numerous challenges (Danuser, 2009; Vallotton and Small, 2009). Fluorescent speckles, sub-resolution clusters of fluorophores (Waterman-Storer and Salmon, 1998), are often faint and unstable, and typically present in only a few images of a time-lapse sequence (Yang et al., 2007). Their motions can be entangled and fast-evolving, especially in complex cytoskeletal structures where large numbers of speckles move in an organized fashion in multiple directions as a part of overlapping speckle flows. Recently, we proposed a new method that we abbreviate as Instantaneous Flow Tracking Algorithm (IFTA), based on graph theory, which can be used to establish motion correspondence between speckles in time-lapse image sequences (Matov et al., 2011). Specifically, IFTA computes multiple vector flow fields based on knowledge of the regional organization and local smoothness of speckle motion (Matov et al., 2011) by considering three consecutive time-frames (Vallotton et al., 2003; Vallotton et al., 2017) of an image sequence at a time. Thus, the method allows us to consider not only the displacement of speckle cohorts, but also the local curvature and acceleration of their motion. This leads to a very accurate assignment of the correspondence between speckles in consecutive frames.

Additionally, we apply bi-objective optimization (Matov et al., 2011) in which the maximization of the number of accepted links between speckles competes with the global cost (computational penalty) of linking speckles. During building the graph solution, when candidate matches form unusual triplets, because of the direction or speed of motion, these links are assigned a higher cost and thus become more likely to be excluded during the overall selection. Naturally, the two goals of forming as many links as possible at the lowest cost are contradictory, as maximizing the number of links requires an increase in overall cost, while minimizing the overall cost requires a decrease in the number of links. This is a key novelty in our approach that guarantees, mostly critically, the elimination of any false positive links.

The generation of false positive tracks has been the most important flaw of all previous tracking approaches (Genovesio et al., 2006; Sbalzarini and Koumoutsakos, 2005; Shafique and Shah, 2005; Vallotton et al., 2003; Vallotton et al., 2017; Veenman et al., 2001) used in the flow tracking scenarios we consider. It creates the issue of delivering wrong flow information and, this way, leads to conclusions regarding cell behavior which do not correspond to the actual processes in living cells. The contribution of IFTA consists of its unique ability to identify organized motion of cohorts of speckles, while removing from consideration the speckles which do not belong to a specific cohort. That is to say that, for instance, if we measure a cell with four flows which move in different directions and there is a fifth group of speckles which undergo unstructured motion, IFTA will dissect the behavior of these four speckle flows, while leaving out the fifth group. The way our solver is built, IFTA only incorporates in its measurements speckles belonging to a cohort, assuming the speckles move in a similar and coherent way, that is, for instance, in the same direction and with the same speed (or acceleration). The cut off values for accepting a speckle within a flow are assigned dynamically based on a Bayesian rule and the posterior distributions of each of the components (weights) of the cost function, which are updated at every step of an iterative procedure.

For visualization purposes, we have applied wide directional averaging (Geusebroek et al., 2003) of the displayed in the figures motion vectors in order to observe and compute the overall differences in the angular orientation of the flow fields, which is another novelty. This filtering comes at the expense of smoothing the local curvature, which is not so clearly observed in the images presented; however, this visualization parameter can be easily modified to show better both the local speed, acceleration, and directionality. Our objective has been to convincingly demonstrate that there are unappreciated cellular overlapped polymer flows that have not thus far been detected and measured. The histograms in all figures are, nevertheless, based on the unfiltered, raw vectors obtained from the clustering analysis after splitting the vectors resulting from the bi-objective optimization into cohorts and therefore represent the true flux speeds for each of the tracked vectors.

IFTA does not overlink speckles and that leads to no false positive selections, while, by avoiding too stringent selections, we minimize the false negative selections too. Thus, finding the optimal balance between the two optimization goals yields a robust tracking of contractile structures with an overall minimal number of false positives and false negatives at the same time. In contrast, the existing specialized speckle tracking approaches are not well suited for the analysis of contractile structures (Ji and Danuser, 2005; Ponti et al., 2005; Yang et al., 2005). Cellular contraction occurs when anti-parallel microtubule (MT) bundles or filamentous actin (F-actin) branches lengthen or shorten in interdigitated polymer meshworks (or networks) that are interdigitated by molecular motors. Hence, cellular structures of the key characteristics studied in this paper and described below generate image datasets exhibiting frequent change of flux speed and directionality on a very short time scale. In principle, there is no limit to how many flows of different directionality can be analyzed with our method. In addition, the cost function and its weights are self-adaptive, and IFTA requires no *a priori* knowledge or user input in regard to motion velocity.

In all examples shown here, earlier tracking approaches have been unsuccessful in resolving in a fully automated fashion the anti-parallel flux of interdigitated cytoskeletal networks when cohorts of speckles constitute coherent flows that overlap. These approaches also fail when confronted with speckle flows that exhibit twist or curl, a situation encountered when cytoskeletal networks flow around fixed objects within the cell or when the activity of molecular motor proteins is inhibited. Therefore, we propose using IFTA as a quantitative tool for the analysis of sub-cellular areas with anti-parallel and overlapping filamentous polymer networks that is capable of precisely measuring the instantaneous change of flow directionality and rapid increase or decrease of the velocity of MT and F-actin flux. We consider the strengths of IFTA to be of particular importance for the evaluation of novel cancer and neurodegeneration treatment compounds. Such drugs modulate the cytoskeleton, sometimes in unexpected ways, and the ability to reverse impaired cellular functionality in disease pre-determines their efficacy in clinical trials. Failure to rigorously probe novel compounds in functional, cellular assays has been the reason for poor lead candidates secondary screening, which, in turn, leads to low cure rates in the clinic (Amiri-Kordestani and Fojo, 2012). Our observation is that this failure is not the result of bad experimental approaches, but rather errors in thoroughly quantifying the available imaging data. These errors stem largely from a lack of specialized computer vision tools required to dissect the incredibly complicated scenarios encountered in the biology of the cell in disease and during treatment in particular.

The main discoveries presented in the manuscript, based on the quantitative analysis of published and newly generated imaging data, are related to the ability of the cell to adapt to a wide range of biochemical and mechanical perturbations. We have investigated this in the context of the two main cytoskeletal proteins, tubulin and actin. Tubulin has been the target of numerous therapeutic approaches for six decades now (Wani et al., 1971; Yang and Horwitz, 2017). Most of these therapies have been, in essence, a “black box” approach (Fuchs and Johnson, 1978; Komlodi-Pasztor et al., 2012; Weaver, 2015), where a treatment with debilitating side effects is administered without having the ability to anticipate whether it would affect the desired target (Kitagawa, 2011; Komlodi-Pasztor et al., 2011; Mitchison, 2012). In this manuscript, a quantitative evaluation of some of the more difficult to interpret effects of tubulin-targeting compounds has been reported together with an attempt for mechanistic explanations for the reasons behind the measured differences. Next, actin is instrumental in wound healing as well as during tumor metastasis; it is also the most important protein in heart and other muscle function. Very few drugs currently modulate actin activity directly, albeit the abundant availability of anti-inflammatory medications that affect the COX pathway (Flower, 2003).

It is the variability of patient response in genomics precision medicine that underscores the importance of utilizing quantitative imaging techniques in the clinic. A plethora of the cellular proteins exhibit ambiguity in function and response to drug perturbations. There exists a high level of regional variability in the dynamics of polymer networks in living cells, which are considerably more interdigitated than initially perceived and reported in the literature. Drugs bind to proteins within these networks and affect their function. Consequently, the modulation of MT and F-actin dynamics and contractility inevitably affects drug efficacy. Answers about the potential efficacy of a drug can, thus, be obtained by developing software systems that measure, in great detail, the effects of a regimen on its cellular targets. The quantitative analysis of molecular manipulations of living patient-derived cells *ex vivo* can reveal which secondary mechanisms would be activated after a particular intervention. Such clinical research can aid the discovery of an optimal drug selection for each disease, which is the motivation for this work.

## MATERIALS AND METHODS

### Sources of FSM datasets

Most of the speckle image datasets presented in this paper were generated previously and computationally analyzed with earlier tracking methods in the context of several publications (Table 1). The new speckle imaging datasets analyzed in this work are the tropomyosin perturbation data and these datasets exemplified in a compelling way the specific strength of our algorithm.

**Table 1.** Data description. This paper presents additional computational analysis of previously analyzed and published image datasets in six manuscripts and two new conditions, two tropomyosin isoforms overexpression, prepared specifically for this publication. IFTA adds a new level of insight in addition to the already available information about speckle flows delivered by earlier tracking methods (Ji and Danuser, 2005; Ponti et al., 2005; Yang et al., 2005).

| **Cytoskeletal**<br>**Filament** | **Condition and Perturbation** | **Sampling**<br>**Rate (sec)** | **Pixel Size**<br>**(nm)** | **Reference** |
| --- | --- | --- | --- | --- |
| **Opposite MT**<br>**Bundles in**<br>**Mitotic**<br>**Spindles** | 1. Wild type | 10 | 67 | (Cameron et al., 2006) |
|  | 2. AMP-PNP, 5 mM | 5 | 67 | (Miyamoto et al., 2004) |
|  | 3. S-trityl-L-cysteine, 100 mM | 5 | 67 | (Miyamoto et al., 2004) |
|  | 4. Monastrol, 0.5–1 mg/mL | 5 | 67 | (Miyamoto et al., 2004) |
|  | 5. Needle skewering<br>*(Xenopus laevis* egg extracts) | 5 | 129 | (Gatlin et al., 2010) |
| **Contractile**<br>**F-Actin**<br>**Meshworks**<br>**in Migrating**<br>**Epithelial**<br>**Cells** | 6. Rac1 activation (see (Matov, 2025a) for analysis) | 5 | 67 | (Wittmann et al., 2003) |
|  | 7. Wild type<br>(PtK1 cells) | 10 | 67 | (Gupton and Waterman-Storer, 2006) |
|  | 8. Wild type | 10 | 133 | (Burnette et al., 2007) |
| | 9. Blebbistatin, 70 μM<br>(Primary cultures of<br>*Aplysia bag* cell neurons) | 10 | 133 | (Burnette et al., 2007) |
|  | 10. Tm2 overexpression | 10 | 67 | New dataset |
|  | 11. Tm4 overexpression<br>(PtK1 cells) | 10 | 67 | New dataset |

### Image analysis

All image analysis programs for detection and tracking of speckles and graphical representation of the results were developed in Matlab and C/C++. A speckle detection (and a MT plus-end tracking) method ClusterTrack used is described and validated in (Matov et al., 2010), the speckle tracking method IFTA is described and validated in (Matov et al., 2011). The computer code is available for download at: https://www.github.com/amatov/InstantaneousFlowTracker. The CPLEX optimizer requires an ILOG license from IBM.

To identify speckles among the numerous bright areas that are convolved with imaging noise, we are mindful that the image signal frequencies are limited by the optical point spread function (Airy, 1835). For analysis, we retains features collected with a band-pass filter with a high spatial frequencies cut off reflective of the microscope optics to suppress noise components (Matov et al., 2010; Ponti et al., 2005) and further exclude non-specific background fluorescent labeling associated with low frequency noise components. The filtering and feature selection are complemented with least squares fitting of an *a priori* unknown number of speckles in the raw image against an average to remove outliers based on shape and intensity (Matov, 2024b). The software performs an automated extraction of speckle tracks and subsequent statistical analysis of the data. The confounding problem in MT and F-actin flux images is the anti-parallel speckle flows combined with a high speckle proximity. We use graph theory - optimizing the network flow through the graph while minimizing the cost of linking - to obtain tracks forming coherent speckle flows (Matov, 2024b; Matov et al., 2011). This approach brings two advantages: (i) speckles can be tracked subject to certain (imposed by a self-adaptable cost function) rules on motion smoothness, resulting from the particularities of MT and F-actin mechanics; (ii) speckles can be tracked under variable topological conditions, that is, in the case of sudden speckle appearance and disappearance, instantaneous change of flux direction or changes in the number of speckle flows (Gatlin et al., 2010). Our approach allows the evaluation of various micro-molecular and pharmacological treatments as well as the testing of specific biological hypotheses (Matov, 2024h).

Extensive calibration and validation has been done of all tracking software modules by (i) manual benchmarking, (ii) benchmarking with known pharmacological perturbations, and (iii) artificial (synthetic) datasets (see in Fig. S1 examples of tracking in artificial datasets representing three-directional motion and circular motion). To compare automated speckle tracking with the gold standard of image analysis and interpretation, manual tracking, we developed a “manual tracker” (Matov, 2024g) module which has a graphical user interface for user navigation. It allows manual tracking of speckles from appearance to disappearance by clicking on the computer screen while the program is saving the lists of selected image coordinates. The users click on speckles in each frame of a time-lapse sequence, linking them to those in the previous. The software stores all data and compares it to the automated tracking results. To validate our results, we also use image sequences of wild type cells and cells subjected to drugs and antibodies perturbing polymer dynamics with known effects. Some of these perturbations, for instance in the context of measuring slower flux and stopping flux by AMP-PNP (Brady, 1985) or monastrol (Mayer et al., 1999), which inhibit motor protein activity, and blocking dynamic instability by paclitaxel (Jordan and Wilson, 2004). Each software module is further benchmarked using artificial (or synthetic) image datasets for which the ground truth is known and the level of tracking difficulty can be modulated during data generation.

In brief, we proposed a new method for instantaneous flow tracking based on network flow theory that can be used to establish motion correspondence between moving speckles in time-lapse image sequences (Matov et al., 2011). The strength of our algorithm is the ability to detect spatiotemporal patterns in organized cohorts of speckles within a larger cohort with unstructured speckle motion. We build triplets based on a search area for each speckle, that is, the maximal displacement computation is limited based on *a priori* information of the video analyzed and utilize a cost function modified from the approach Sethi proposed (Sethi and Jain, 1987), and compute the Mahalanobis distance (Mahalanobis, 1936). Each detected speckle is initially linked in multiple candidate triplets – in the case of the mitotic spindle, where the speckle density is high, this can be up to 50 triplets. We then use linear programming (Murty, 1985) optimization to select the links that maximize the flow through the graph while minimizing the cost of linking by solving the primal simplex method (Dantzig, 1987). There exist prior algorithms for the tracking of coherent motion (Zhou et al., 2012) in the context of the analysis of multi-directional and anti-parallel flows, which, however, are not interdigitated. The difficulty in solving the motion patterns when the speckles of the different cohorts interdigitate and intermingle is that the distance traveled by the speckles between two time-frames can oftentimes be significantly higher than the average distance to their nearest neighbor. Such scenarios often lead to wrong, false positive assignments. For these reasons, we employ local and global motion models, which are parameters-free and data-driven, to ensure that links from one time-frame to the next are established only when there is evidence for organized motion within a cohort. We rely on a Bayesian statistics approach to fine-tune the weights of the objective function (Matov et al., 2011) and the solution converges (<5% of the links change) in three to four iterations without requiring user parameter input. To implement the angular components of our motion models, a distinct feature of our approach is that it always considers three time-frames at once (Vallotton et al., 2003; Vallotton et al., 2017), that is, we always analyze triplets of images.

This approach allows us to study the distribution of the tentative angular motion of individual speckles in comparison to the overall directions of motion of the identified cohorts. In this context, our cost function for linking speckles is built of three elements, which penalize links that are not equidistant in the steps from frame 1 to frame 2 and from frame 2 to frame 3, also links that display motion with very sharp turns as well as links that do not conform with the overall speed and motion direction of the overall cohort. Following a circular *k*-means clustering (Matov et al., 2011), we apply spatial regularization and the vectors belonging to each of the cohorts (flows) are filtered with an anisotropic Gaussian (Weierstrass, 1885) kernel with a large standard deviation along the flow direction (σ_1_) and a small standard deviation perpendicular to the flow direction (σ_2_). A typical ratio σ_1_/σ_2_ that we use in our analysis is 4, for instance (σ_1_, σ_2_) = (12, 3) or (16, 4), depending on the application. Our global motion model is calculated based on the angle each vector forms with the associated flow vector, that is, it reflects how much a vector is affected by the directional smoothing. This way, we inherently favor linking speckles that move together in spatiotemporal patterns, including accelerating and/or curve patterns, and can be assigned with a minimal penalty to one of the cohorts (Matov et al., 2011).

For image sequences in which a mitotic spindle moves, image registration was done based on cross-correlation analysis during which the translation was corrected via evaluating the covariance matrix by computing the convolution of image pairs (Mitchison et al., 2004b). Comparisons between the tracking results of IFTA and the feature point tracking algorithms developed in (Genovesio et al., 2006; Sbalzarini and Koumoutsakos, 2005) are presented in (Matov, 2024b). One of our motion models (the global motion model), penalizes the deviation in terms of angular orientation of the raw vectors from the estimated overall flow direction; it is calculated based on the angles between the raw vectors and the computed flow field vectors. The flow fields are obtained by clustering with *k*-means (Steinhaus, 1957) of the raw vectors, which allows vector assignment to different populations of moving speckles. Further, most tracking algorithms maximize the number of links (Shafique and Shah, 2005; Vallotton et al., 2003; Veenman et al., 2001), thus introducing false positive links, for example, when tracking cohorts of speckles that do not have a constant number of participants from one time-frame to the next, because of occlusions. To improve on that strategy, we introduced graph pruning. In this context, the most important feature of our approach is the application of bi-objective optimization. We use Pareto efficiency (Pareto, 1897), a method widely used in microeconomics, which Vilfredo Pareto conceived in the 19^th^ century, after realizing that two concurrent aims could be competing by definition and the most optimal solution is the point on the convex curve, which is closest to zero, the so-called utopia point. This way, we compute the optimal point of a trade-off between forming as many links as possible on one hand - which inherently increases the overall cost (or penalty) - and minimizing the cost of linking on another. That is to say that by posing competing aims to our solver, our objective is to only select low cost links. This way, we reduce to an absolute minimum the false positive selections we make, which is of critical importance. The weights of the Pareto optimality multi-objective function are based on Bayesian statistics, which makes the algorithm self-adaptive with rapid convergence within several iterations (most commonly, within three to four iterations less than 1% of the links change between selection steps). During the analysis of mitotic spindles, the optimization of the cost function weights further converged after five iterations, when only 0.5% (Matov, 2024b) of the triplets changed in comparison to the previous iteration.

We evaluated the performance of another tracking algorithm, which uses three different motion priors, models of Brownian motion and directed movement with constant speed or acceleration, using a maximum likelihood association (Genovesio et al., 2006). The software can track a total number of 20,000 trajectories and the test mitotic spindle image sequence contains about 50,000 speckles that form short trajectories and new speckles continuously appear throughout the time-lapse sequence. This scenario represents a tracking challenge and the software did not track a significant proportion of the speckles. In addition, the trajectories often represented nearest neighbor tracking and linked features laterally, which is erroneous, as all features move vertically in one direction only and form spatiotemporal patterns, that is, either roughly half of the speckles move only upwards and the other half move only downwards (Matov, 2024b).

In the simple scenario of two cluster centers (such as the spindle poles) from where vectors originate in each direction radially, we developed a circular *k*-means algorithm (Matov et al., 2011), which allows us to assign each vector to a cluster and then re-calculate the cost values based on the updated flow field vectors. In scenarios in which we are not sure of the number of overlapped flow fields, we utilized a maximization-optimization algorithm (Figueiredo and Jain, 2002) to identify the number of cohorts moving in different directions at the same time. For instance, if we analyze a multi-polar spindle (Matov et al., 2011) and there are several groups of speckles moving toward each of the poles and these groups intersect in the middle of the cell, we will track the exact number, location, speed and direction of motion for each moving speckle and will assign the speckle to the corresponding cohort (Matov, 2025b). The algorithm is not limited in the number of cohorts it can detect and track. To compute circular expectation-maximization, the assignment uses a mixture of Tikhonov/von Mises distributions (Bouberima et al., 2010).

Scale-space theory was popularized during the 90s through the work of Tony Lindeberg (Lindeberg, 1993). It explores, through a cascade of different levels of Gaussian smoothing (Weierstrass, 1885) of an image, the changes in perception depending on the scale of observation. Our initial speckle detection kernel was a difference of Gaussians (DoG) bandpass filter (Matov et al., 2011; Wilson and Giese, 1977). The upper spatial frequency cut-off, reflecting the limit of the optical transfer function (Fourier, 1822) beyond which the collected signal is white noise, was computationally fitted to the Bessel function forming the point-spread function of the imaging system (Bernoulli, 1728), that is, matched to the optical resolution limit of the collection frequency spectrum based on the dye emission wavelength and the diffraction limit of the objective. The lower spatial frequency limit relates to the computational clipping of image background, that is, the non-specific fluorescent signal forming larger shapes in the body of the cell and the cut-off was empirically obtained from the available imaging datasets to minimize the false positive feature selections. The ratio of the standard deviation of the two 2D Gaussian functions we obtained was 1.1 (Matov et al., 2011). David Lowe performed a similar analysis for the scale-invariant feature transform (SIFT) and obtained a ratio value of about 1.4 (Lowe, 1999).

The DoG detection approach allowed us to select statistically significant speckles with high precision at a very low computational cost. When the images contain features that are not diffraction-limited in size, such as the end binding protein 1 (EB1) comets forming at the tips of polymerizing MTs, the feature selection can be based also on detectors such as SIFT (Lowe, 1999; Matov, 2025b), SURF (Bay et al., 2008; Matov, 2024i), or ORB (Rublee et al., 2011). The strength of these algorithms is that they detect features that persist along multiple scales of observation and can be reliably tracked in time, even in scenarios with different lighting and partial occlusion. Having multiple features detected for the same object will also facilitate our flow tracking step since it will generate several vectors with a very similar cost (Matov, 2024b). In the case of tracking motion of EB1 comets, the bag-of-bags-of-words model (Ren et al., 2013) can be applied to the analysis of individual images in order to add an additional component of the cost function. It offers an improvement to the bag-of-words model, because of the inclusion of image spatial information during classification.

Our algorithm uses a global motion model that relies on directional averaging of vectors in order to compute the cost to assigning a link to a cohort based on the vector immediate environment (Geusebroek et al., 2003). When the image intensity is sparse, that is, the speckle density is low and in a certain area there are not many speckles to interpolate over, the flow estimation may fail in determining the number of cohorts or the dominant flow direction. To circumvent this potential issue, in the original clustering implementation we utilized a sliding window approach in which vectors from five image triplets (for instance, from time-frame triplets 1-2-3, 2-3-4, 3-4-5, 4-5-6, and 5-6-7) were aggregated at each time step. This approach always generated the correct flow direction results, but it slowed down the performance of the tracker. To improve on this strategy, we have replaced it with the utilization of Markov Random Field (MRF), which runs as an iterative process over the solution for adjacent image triplet flow fields. It modifies the selection results to reflect consistency with the rest of the flow directions and imposes constraints in order to exclude flow direction selections which are outliers. This way, our existing algorithm has been improved by optimizing the objective function and applying MRF (Boykov et al., 2001) to resolve the topology in sparse datasets without having to aggregate multiple image triplets to resolve the flow directions. For additional examples of tracking results obtained with IFTA, please see (Matov, 2024g; Matov, 2024h; Matov, 2025a; Matov, 2025b).

We have developed ClusterTrack (Matov et al., 2010), a computational approach for the analysis of MT dynamics in living cells by automated tracking of the motion of MT ends in high-resolution confocal microscopy image sequences using a fluorescently tagged MT EB1 comets (Su et al., 1995). EB1s recognize conformational changes and bind the MT tips during polymerization (growth) (Tirnauer and Bierer, 2000). When EB1s are EGPF-labeled, the EGFP-EB1 dimers at the MT tip form a fluorescent comet, which serves as a molecular marker for the direct visualization of MT polymerization dynamics. Our image analysis algorithm tracks thousands of EB1-EGFP comets visible in an image time-lapse sequence allowing the detection of spatial patterns of MT dynamics and introduces spatiotemporal clustering of EB1-EGFP forward tracks to infer MT behaviors during phases of pause and shortening when the MTs are invisible (Matov et al., 2010). EB1 comet detection is accomplished by band-pass filtering, which enhances image features of the expected size of an EB1 comet while it oppresses higher-frequency noise and lower-frequency structures representing larger aggregates of fluorescent protein. The algorithm is robust against variations in comet lengths, both within videos and between videos. The feature detector delivers not only the position of each comet in a time-frame but also the eccentricity and angular orientation of the comets. The latter is used as a directional cue in the subsequent tracking of comets from one time-frame to the next (Yang et al., 2005). The cost of an individual assignment is defined in statistical terms as the Mahalanobis distance (Mahalanobis, 1936) between projected and extracted comets. Because of the lack of direct molecular markers of MT pausing or depolymerization, MTs become invisible when the MTs are not polymerizing. Our algorithm allows us to infer the stages of pausing and depolymerizing by computational clustering of visible polymerization (forward) tracks. Clustering of collinear polymerization tracks that likely belong to the same MT is performed by solving a linear assignment problem (LAP) (Jonker and Volgenant, 1987), which identifies the globally optimal positional correspondences between the endpoints of terminating tracks and the start points of newly initiated EB1 tracks, and every potential assignment is characterized by a cost of linking. A pairing between a track termination and a track initiation is considered only if these two events are separated by less than 15 seconds or another use-specified time-lag. Furthermore, the track initiation has to fall within either one of two cone shaped, geometrical search regions. The choice of the forward cone with an opening half-angle of 60° is motivated by the observation that during pauses or out of focus movements, MTs sometimes undergo significant lateral displacements and directional changes, especially at the cell periphery. The longer the gap (or the distance between visible tracks, see Fig. 1) duration, the larger is the probability for a lateral step and directional change. The reasons for the gaps can also be in and out of focus motion of the MTs, that is, when a MT temporarily leaves the focal plane, or the delay in signal formation as the EB1 comets take a short period of time to form at the polymerizing MT tips as (even if at some instances one can see the outline of the MT) the automated detection algorithm segments only fully formed comets (see examples in Vid. 1 and Vid. 2). In contrast, the backward cone has an opening half-angle of 10° only, motivated by the observation that MT rescues generally follow the polymerization track before the catastrophe. By construction, this assignment is spatially and temporally global, which warrants a high level of robustness. Competing pairings are weighed against one another via a cost function that prefers collinear track pairings with a short gap distance. For each condition, several image sequences containing one or two cells are analyzed. For all conditions, statistical analysis for every MT dynamics parameter is performed using Kolmogorov-Smirnov statistics (Kolmogorov, 1933).

**Figure 1.**
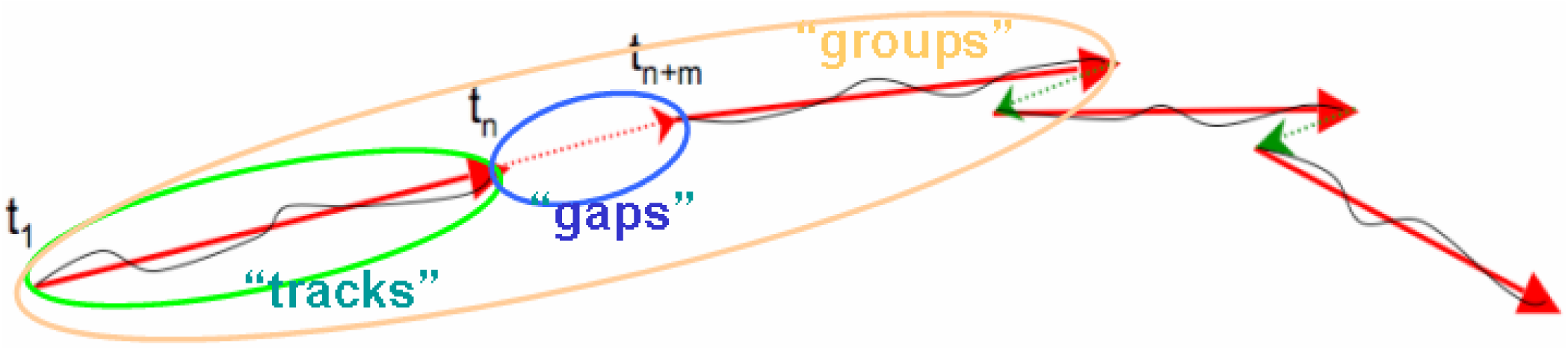
Clustering of EB1 comet trajectories. ClusterTrack (Matov et al., 2010) computationally groups multiple EB1 trajectories (shown in black). The visible (forward) tracks are shown in red (head-to-tail, with arrowheads denoting the direction of motion). Pairing between visible polymerization tracks is performed based on co-linearity and limited by velocity considerations. Gaps (punctuated red line between time-frame n and n+m) between EB1 tracks represent stages of MT pausing and molecular motor flux. Groups represent periods of consecutive polymerization (tracks) and pausing/flux (gaps), which are not interrupted by a catastrophe event, that is, they persist until a switch to depolymerization occurs. The invisible stages of depolymerization are shown in punctuated green lines with arrowheads denoting the direction of motion.

### Cell culture

Kangaroo rat kidney epithelial PtK1 cells (gift from the Fowler lab) were cultured as described previously (Wittmann et al., 2003).

Human Tm2 and Tm4 sequences, derived from the NCBI GenBank database, were cloned from MDA-MB-231 receptor triple-negative breast cancer cells. Total RNA was isolated using RNeasy Mini Kit (Qiagen), then retrotranscribed, and the specific tropomyosin sequences were amplified by PCR.

The long (Tm2) and short (Tm4) human isoforms were ligated into the pIRES2-AcGFP1 expression vector (Clontech). PtK1 cells were transfected with Nucleofector™ Technology (Lonza) with recombinant pIRES2-AcGFP1 vectors expressing Tm2 and Tm4.

X-rhodamine–conjugated actin was prepared as described previously (Waterman-Storer, 2002) and microinjected into cells at 1 mg/mL.

Organoids with diameter 40-80 µm were plated in 30 µL Matrigel drops in six wells per condition of a 24-well plate per condition and treated for two or more weeks with docetaxel with concentration of 10 nM, 100 nM, and 1 µM to eliminate the vast majority of the organoid cells and induce a persister state in the residual cancer cells. This treatment was followed by co-treatment with docetaxel 100 nM and the small molecule ML210 5 µM or docetaxel 100 nM combined with the small molecule ML210 5 µM and lipophilic antioxidants ferrostatin-1 2 µM, rescue treatment. Organoids with diameter 40-80 µm were plated in 30 µL Matrigel drops in six wells per condition of a 24-well plate and also treated for 2 or more weeks with FOLFOX (folinic acid, fluorouracil, and oxaliplatin) with concentration of 1 µM, 10 µM, and 100 µM to eliminate the vast majority of the organoid cells and induce a persister state in the residual cancer cells.

Primary and retroperitoneal lymph node metastatic prostate tumor and sternum metastatic rectal tumor tissues were dissociated to single cells using modified protocols from the Witte lab. Organoids were seeded as single cells in three 30 µL Matrigel drops in 6-well plates. Organoid medium was prepared according to modified protocols from the Clevers lab and the Chen lab.

To obtain a single cell suspension, tissues were mechanically disrupted and digested with 5 mg/mL collagenase in advanced DMEM/F12 tissue culture medium for several hours (between 2 and 12 hours, depending on the biopsy or resection performed). If this step yielded too much contamination with non-epithelial cells, for instance during processing of primary prostate tumors, the protocol incorporated additional washes and red blood cell lysis (Goldstein et al., 2011). Single cells were then counted using a hemocytometer to estimate the number of tumor cells in the sample, and seeded in growth factor-reduced Matrigel drops overlaid with prostate cancer (PC) medium (Gao et al., 2014). With radical prostatectomy specimens, we had good success with seeding 3,000 cells per 30 μL Matrigel drop, but for metastatic samples organoid seeding could reliably be accomplished with significantly fewer cells, in the hundreds.

### Sample processing

Small RNA were extracted from de-identified urine samples (50 mL) by Norgen Biotek Corp.

Retroperitoneal lymph node organoid whole genome sequencing (736 ng DNA) was done at the UCSF Core Facilities.

### Microscopy imaging

Cells were maintained on the microscope stage at 37°C with an air stream incubator (Nevtek) in aluminum chambers (Wittmann et al., 2003) in culture medium containing 30 µL oxyrase per milliliter of media (Oxyrase, Inc.) to inhibit photobleaching. F-actin FSM time-lapse image series were acquired at 10 seconds intervals using a 100x 1.49 NA Plan Apo TIRF DIC objective lens (Nikon) on a spinning disk confocal scanner (Yokogawa) as described in (Adams et al., 2003). F-actin FSM time-lapse image series were acquired on an inverted microscope (model TE2000, Nikon) equipped with electronically controlled shutters and a robotic stage with linear position feedback encoders on the x, y, and z axis (model MS-2000, Applied Scientific Instruments). Images were acquired on a 12-bit cooled CCD camera (model Coolsnap HQ^2^) controlled by MetaMorph software (Universal Imaging Corp.).

Organoids were imaged using transmitted light microscopy at 4x magnification and phase contrast microscopy at 20x magnification on a Nikon Eclipse Ti system with camera Photometrics CoolSnap HQ2.

Organoid long-term imaging was done over 72 hours by taking an image every 10 minutes at 20x magnification.

We imaged EB1ΔC-2xEGFP-expressing organoids (Koo et al., 2013) by time-lapse spinning disk confocal microscopy using a long working distance 60x magnification water immersion 1.45 NA objective. We acquired images every half a second for a minute (that is, 120 images) to collect datasets without photobleaching (Gierke and Wittmann, 2012).

## RESULTS

### Tracking spindle MT movements

MTs constitute the major structural element of the mitotic spindle (Inoué and Ritter, 1975). Within this bipolar, fusiform structure, inherently polar MTs are typically oriented with their minus ends, from which tubulin dimers dissociate, pointing toward one of the two poles and their plus ends, where tubulin dimers are added to the MT lattice and begin moving toward one of the poles, pointing toward the midzone (Sharp et al., 2000). This spatial organization creates a region of predominantly anti-parallel MT overlap within the central part of a bipolar spindle where MT-based motor proteins, such as heterotetrameric kinesin-5 (Eg5 in *Xenopus*, KIF11 in human), cross-link and slide MTs resulting in slow poleward transport of MTs (Kapitein et al., 2008; Kapoor and Mitchison, 2001). Termed “flux” (Dietz, 1972; Inoué and Salmon, 1995), the abovementioned movements can be visualized by FSM (Waterman-Storer et al., 1998). The anti-parallel flux in the central spindle represents a difficult, yet well-characterized tracking problem. There are important subtleties (Vallotton et al., 2003), however, in the organization of the spindle, such as the slower flux in kinetochore MTs compared to non-kinetochore MTs, which remain largely unexamined. Statistical analysis of large populations of speckles can deliver insights regarding the differences in the speckle distributions in question when the underlying analysis extracts the correct data vectors. When the data is analyzed without the necessary motion models and optimization techniques, false positive links lead to the incorporation of unexisting motion components and that, in turn, leads to inability to distinguish faint differences between different flow distributions. Reporting such differences becomes very important in disease and during patient treatment as novel regimens affect differentially the variety of MT subpopulations depending on their function and regulation.

It becomes even more challenging for the traditional techniques to measure flows in multipolar spindles (Therman and Timonen, 1950) in cancer cells. Spindles which progress to anaphase without forming a bipolar structure result in aneuploidy. IFTA was successfully tested to resolve the flows in a tri-polar spindle in a *Xenopus* egg extract (Matov et al., 2011), which was accidentally formed when several MT bundles had glued to the cover slip, thus forming artificially a third pole. Our algorithm can resolve any number of overlapped flows (we perform tests for the existence of between one and nine flows) (Matov, 2024b; Matov, 2025b), which can be valuable when testing the effects of novel therapies to, for instance, affect multipolar spindles selectively.

#### Anti-parallel MT flux in wild type mitotic spindles

As an initial test of IFTA, we used our algorithm to measure MT flux rates in “cycled” spindles assembled in *Xenopus* egg extracts (Desai et al., 1999), some of which form about 3,000 speckles at each time-frame. IFTA was readily able to detect opposing speckle flows in the spindle midzone. Figure 2A shows a single FSM image of labeled-tubulin speckles from an FSM time series (untreated spindle image sequences were taken from (Cameron et al., 2006)) that has been overlaid with flow vectors generated using IFTA. Two opposing flows were detected and clustered based on their distinct directionalities toward each of the two spindle poles (shown in red and yellow vectors). Note the large spatial overlap, particularly in the spindle midzone, between the opposing flow fields. The main flux directions detected are marked with a red arrow, toward the right pole, and a yellow arrow, toward the left pole. The IFTA-measured flux velocities (Matov and Danuser, 2004) were similar for several FSM datasets acquired using either spinning disk confocal fluorescence microscopy or widefield microscopy (2.02 ± 0.73 µm/min and 2.21 ± 0.81 µm/min, respectively) and comparable – albeit 12% slower - to published rates measured using linear Kalman filter tracking (Yang et al., 2005) in widefield microscopy FSM images (Yang et al., 2008) (2.52 ± 0.65 µm/min). Cross-correlation tracking in widefield microscopy FSM images (Miyamoto et al., 2004) (1.97 ± 0.16 µm/min), by design of the methodology, underestimates the flux rates, while the kymograph analysis (Miyamoto et al., 2004) (2.21 ± 0.45 μm/min) is consistent with our measurements with IFTA. Thus, even though IFTA represents a fundamentally different tracking strategy, these data demonstrate that the approach can both detect and accurately measure spindle MT flows.

**Figure 2.**
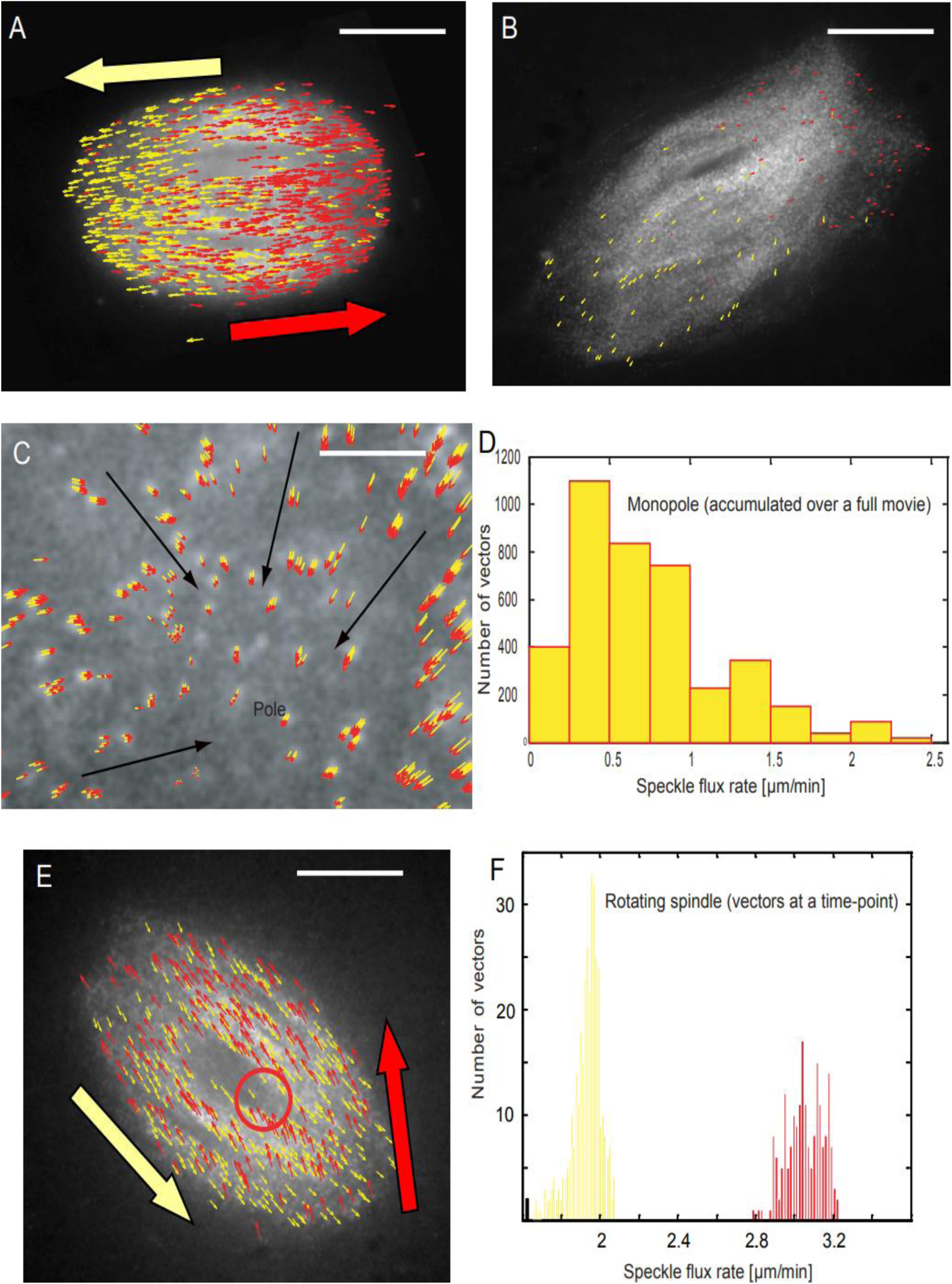
IFTA analysis of spindle MT dynamics in *Xenopus laevis* egg extracts. IFTA was used to track bi-directional MT polymer flux in metaphase spindles in *Xenopus* egg extracts and imaged using FSM. Figure 2A. The figure shows a single FSM image of labeled-tubulin speckles from an FSM time series (untreated spindle image datasets were taken from (Cameron et al., 2006)) that has been overlaid with flow vectors generated using IFTA. Two opposing flows were detected and clustered based on their distinct directionalities toward each of the two spindle poles (shown in red and yellow vectors). The main flux directions detected are marked with a red arrow, toward the right pole, and a yellow arrow, toward the left pole. Scale bar equals 5 µm. Figure 2B. Bipolar spindles were imaged using FSM following the addition of 5 mM AMP-PNP (a non-hydrolysable ATP-analog that inhibits MT-based motor activity) to analyze spindle assembly reactions. IFTA, in contrast to earlier analysis methods, measured about 90% inhibition of MT flux. However, a small number of motion vectors (similarly to Fig. 2A, shown in red and yellow to display vector direction of motion toward each of the two spindle poles), fewer than 130 out of over a thousand speckles in each image (about 10%), reported MT movements. The average measured speed is 0.2 µm/min, and the maximal speed is 0.6 µm/min. These localized speckle movements resembled contractions and seemed to buckle MT bundles in proximity to chromosomes. Scale bar equals 5 µm. Figure 2C. The figure shows vector fields superimposed on FSM data of Eg5-inhibited monopoles (image datasets were taken from (Miyamoto et al., 2004)). IFTA measured median flux rates of about 0.6 µm/min in these structures (see also Vid. 4), with the fastest flux speeds being up to 2.7 µm/min (see the histogram in Fig. 2D). Vectors are drawn in black color to mark the inward speckle flow motion toward the center of the pole of the monopolar spindle. The head of all individual speckle vectors are displayed in red to show the inward direction of the speckle motion. Scale bar equals 2 µm. Figure 2E. In the figure, IFTA was used to track speckle flow in spindles skewered with a single, vertical microneedle (image datasets were taken from (Gatlin et al., 2010)). Under these conditions, spindles slide parallel to their interpolar axes and, at the same time, rotate continuously with velocities slightly slower than those typically measured for flux, ultimately escaping the needle entirely. The red circle indicates the needle position and a red arrow depicts speckle motion away from the needle (toward the top of the image, red vectors) whereas a yellow arrow depicts speckle motion toward the needle (that is, toward the lower right corner of the image, yellow vectors). Scale bar equals 5µm. The histogram of flow velocities (Fig. 2F) shows two distinct velocity populations. The red population displays speckle motion away from the needle, with an average speed of over 3 µm/min; in yellow is displayed the significantly slower (with 1.1 µm/min or 37% slower), with an average of just over 1.9 µm/min, speckle motion toward the needle.

#### Incomplete inhibition of MT flux in mitotic spindles after treatment with AMP-PNP

IFTA uses data-driven self-adaptive local and global speckle motion models (Matov et al., 2011). A well-known problem with computational tracking methods which apply motion models is the formation of false positive links between successive imaging time-frames when there is no actual motion. To benchmark IFTA and to test whether it generates false positive links, we analyzed bipolar spindles imaged using FSM in the presence of 5 mM AMP-PNP (Miyamoto et al., 2004), a non-hydrolysable ATP-analog that inhibits MT-based motor activity, which is used in the analysis of spindle assembly reactions. This treatment has previously been reported to completely inhibit spindle MT flux while maintaining a bipolar spindle morphology (Sawin and Mitchison, 1991). Previous kymograph-based analysis (Kapoor and Mitchison, 2001) suggested no MT movement in such spindles already in the presence of 1.5 mM AMP-PNP and certainly no concerted flows. IFTA (as well as the visual inspection of Vid. 3), in contrast, detected an inhibition of MT flux for about 90% of the speckles, but identified a small cohort of moving speckles for which we measured an average speed of 0.2 µm/min and a maximal speed of 0.6 µm/min (Fig. 2B, n = 129 speckles).

Roughly 10% of the speckles in each image exhibited motion and indicated contraction in several of the MT bundles oriented toward the corresponding pole of the spindle (Fig. 2B, the yellow vectors are in one half of the spindle and point toward the closer pole and the red vectors are in the other half of the spindle and point toward the opposite pole). These localized speckle movements and contractions - resembling the way a snail crawls (see also Vid. 3) - seem to buckle the MT bundles around the chromosomes. These results suggest a residual motor activity in these spindles and potent contractions at several locations. This completely novel insight in the effects of AMP-PNP suggests a lower inhibition of cytoplasmic dynein (Gibbons, 1963) at the minus ends of bundled MTs, even in areas of the spindle away from the poles. In *Drosophila* embryos, the inhibition of KLP61F (an Eg5/KIF11 homolog) leads to an 80% reduction in flux speeds and, in some cases, a negative flux, that is, a reverse in the flux direction (Brust-Mascher et al., 2009), which is likely the result of MT depolymerase (Brust-Mascher et al., 2004) or/and dynein activity. Such secondary mechanisms may explain the limits of the potential use of AMP-PNP, and similar agents, like the KIF11 inhibitor ispinesib (Burris et al., 2011), in therapeutic applications. It points toward the need to combine compounds, such as the addition of a dynein inhibitor (Höing et al., 2018), and test their effects on patient-derived cells *ex vivo* before any utilization in a clinical setting.

#### Residual MT flux in Eg5-inhibited monopoles

Small molecule motor inhibitors, like monastrol and S-trityl-L-cysteine, specifically target the flux-generating machinery by affecting the function of Eg5/KIF11. Spindle assembly reactions in Eg5-inhibited extracts produce monopolar MT arrays, whereas the addition of these inhibitors to extracts with pre-assembled bipolar spindles results in spindle collapse into monopolar structures (Mitchison et al., 2004a). Previous analysis of MT flux in Eg5-inhibited monopoles have yielded inconsistent results for the measured average speckle speeds. Image cross-correlation (<0.2 µm/min) and kymograph (0.32 µm/min) analyses suggested insignificant flux rates in Eg5-inhibited spindles (Miyamoto et al., 2004), leading to the conclusion that Eg5/KIF11 motors are the sole drivers of flux in extract spindles.

Conversely, when IFTA was used to measure flux rates in the same monopolar structures, the results indicated a median flux rate of about 0.6 µm/min (Fig. 2C, Fig. 2D, and Vid. 4), with the fastest speeds being up to 2.7 µm/min. In Fig. 2C (speckle flux vectors have read arrowhead to highlight the direction of motion), vectors are drawn in black color to mark the inward speckle flow motion toward the pole of the monopolar spindle. The absence of active Eg5/KIF11 indicates the presence of MT sliding in such monopoles with inhibited MT treadmilling. Our new results indicate that other proteins are contributing to the generation of spindle MT flux - independent of Eg5/KIF11 activity - as the detection of flux in Eg5-inhibited monopoles indicates the presence of additional, Eg5-independent flux mechanisms. This result is consistent with observations in mammalian tissue-culture cells where it was hypothesized that a reeling-in force, generated by spindle pole-associated MT depolymerases, also contributes to flux (Cameron et al., 2006; Gaetz and Kapoor, 2004; Rogers et al., 2005). Furthermore, the detection of flux in monopoles lends credence to spindle assembly models that inherently rely on MT transport to generate spacing between chromosomes and poles (Brugués et al., 2012). Further, flux in Eg5-inhibited monopoles measured using IFTA is similar to that detected in the “polar MT bundles” that remain following treatment with Op18, a MT-destabilizing protein that selectively target the “barrel MT bundles” constituting the bulk of MT mass near the spindle midzone (Houghtaling et al., 2009; Yang et al., 2008). Taken together, these measurements (Matov and Danuser, 2004) suggest, in both abovementioned scenarios, consistent cytoplasmic dynein (Gatlin et al., 2009; Li et al., 2007) activity.

#### Fast-evolving MT flux in mitotic spindle manipulated by microneedles (skewered spindles) exhibiting unpredictably twisting motion with rapid changes in directionality

A particularly well-suited application to demonstrate the unique capabilities of IFTA is the study of structural occlusions in the spindle that locally influence the usual poleward directionality of flux. Artificial inclusions can be introduced by skewering extract spindles with microneedles (Gatlin et al., 2010; Shimamoto and Kapoor, 2012). In previously published work, IFTA was used to make inferences regarding the mechanical properties of the putative spindle matrix (Gatlin et al., 2010). Spindles skewered in this way pushed themselves off the needle by sliding parallel to their interpolar axes and, at the same time, rotated continuously at velocities slightly slower than typical flux rates, ultimately escaped the needle entirely (Gatlin et al., 2010). The red circle indicates the needle position needle in Fig. 2E and red arrows depict speckle motion away from the needle (toward the top of the image) whereas yellow arrows depict speckle motion toward the needle (that is, toward the lower right corner of the image).

These observations suggested that forces generated by flux are transmitted to the needle, hypothetically via MT cross-links that flow with the moving lattice. This means that MTs would flow around the needle at velocities equal to the velocity of flux within the spindle minus the escape velocity, that is, *v̅*_1_ = *v̅*_flux_ − *v̅*_spindle_, (Fig. 2E, yellow arrows pointing to the right and downward), while the velocity of flux in the opposite direction is increased to *v̅*_2_ = *v̅*_flux_ + *v̅*_spindle_ (Fig. 2E, red arrows pointing up, in the direction of rotation; see also the clearly separated speed distributions in Fig. 2F). Similar to a flag in the wind, in these experiments – similarly to mitotic spindles in cancer of the breast and the colon (Matov, 2024f) – rapid, fast-evolving flows in the *Xenopus* egg extract often caused spindles to rotate around the skewering needle.

The histogram of flow velocities (Fig. 2F) shows two distinct velocity populations. The red population displays speckle motion away from the needle, with an average speed of over 3 µm/min; in yellow is displayed the significantly slower (with 1.1 µm/min or 37% slower), slightly over 1.9 µm/min, speckle motion toward the needle. Because the flow vectors rapidly change their directionality near the needle, linear Kalman filtering (Yang et al., 2005) links features across fibers and fails to accurately follow the local speckle motion. We studied nine such spindles and IFTA measured consistent significant differences in the average flow speed depending on flux direction, toward or against the direction of spindle motion (comprised of directional displacement and rotation of one of the poles, most commonly the pole further away from the needle) while escaping the needle. In all spindles the concrete trajectory of overall motion was very different and for all spindles IFTA could accurately follow every step of the rapidly changing spindle location. See Table 1 in Materials and Methods for further information regarding the conditions and perturbations.

Taken together, these new results indicate that IFTA is the algorithm of choice to measure instantaneous fast-evolving overlapped flow fields of cohorts of speckles participating in multi-directional organized motion. Our method also has particular strengths and robustness when the data pose substantial challenges in terms of spatiotemporal variation of speckle behavior within the flow field, as well as speckle appearances and disappearances. IFTA captures the instantaneous organization of the speckle flows by balancing the number of speckles linked with how well those links conform to the overall motion of the speckle flow they are assigned to. The trade-off between the numbers of links formed versus the overall cost (where a lower cost is a measure of similarity) of linking is accomplished by performing bi-objective optimization between the local and regional motion models; it can be imposed adaptively to the continuously changing speckle dynamics. Thus, any re-organization in the overlapping speckle flows is captured and measured by using as few as three consecutive images. Therefore, IFTA will be very suitable for the analysis of MT flux at other very delicate stages of cell division, such as prometaphase and the onset of anaphase. At these stages, the spindle behavior is somewhat similar to that of the skewered spindles described in this section. Namely, the flux in a portion of the MT bundles suddenly accelerates leading to irregular patterns of speckle flow fields and, in some cases, rotational motions and rocking of the spindle. The spindles rapidly change position for a short period of time, in a similarly unpredictable manner, before continuing into the following stage of cell division.

Our novel results demonstrate the extraordinary ability of the mitotic spindle to re-arrange and correct mechanical imperfections in its structure. The quantitative analysis indicates the activity of forces related to the extracellular matrix and molecular motor proteins are driving these re-arrangements, rather than MT polymerization dynamics. The clinical implication of these measurements is that even the maximum tolerated dose of docetaxel, which does not arrest all spindles in mitosis but rather many, would allow for a few tumor cells to proliferate.

As an example, in PC, most tumors are heterogeneous and dormant – until they are treated. After initially receding, the tumors, however, resume growing. Further, the drug treatment process would effectively serve as a repeated selection step, allowing for the replication of the most resilient clones. Taken together, the data indicates the level of difficulty in interfering with mitosis by modulating cytoskeletal polymers without inhibiting the action of specific plus-end and also minus-end directed molecular motors as well as proteins providing mechanotransduction in the tumor microenvironment.

We performed a similar analysis of rapidly rotating mitotic spindles in treatment-resistant receptor triple-negative breast cancer MDA-MB-231 cells (Matov, 2024f). Using IFTA, we determined the number of flows on-the-fly and assigned the tracking vectors to each of the corresponding flow fields. Further, to visualize the angular orientation of speckle flows, we dynamically color-code each flow, that is, the color of each flow changes as the spindle rotates to improve visualization (see an example in Vid. 5).

### Tracking contractile actin flows in fluorescent speckle microscopy data

Similarly to spindle MT arrays, F-actin filaments are often arranged in anti-parallel contractile fibers or in more complicated contractile meshworks, for example near the leading edge (Gov and Gopinathan, 2006; Mitchison and Cramer, 1996; Small et al., 1978) of migrating cells. Cell migration is powered by a coordinated cycle of leading-edge protrusion (Cramer, 1999; Mogilner and Oster, 1996; Oster and Perelson, 1987) in the direction of migration, substrate adhesion of the protrusion, generation of tension on new focal adhesions (FAs) (Kaverina et al., 2002) to advance the cell body (Small and Resch, 2005), and de-adhesion of the trailing cell rear. F-actin is required for each step of the cycle (Welch et al., 1997). Spatio-temporally coordinated regulation of the interaction of F-actin with specific binding proteins and myosin motors is required for the actin cytoskeleton to perform such diverse mechanical functions. Because of a lack of appropriate computer vision methods, the dynamics of contractile F-actin structures has been understudied. We, therefore, proceeded and tested the applicability of IFTA to the investigation of contractile actin network arrangements followed by rapid F-actin meshwork re-arrangements in migrating epithelial cells and similarly highly dynamic neuronal growth cones.

#### Anti-parallel actin flows in the “convergence zone” between lamella and cell body in migrating epithelial cells

Anti-parallel F-actin (Kolega, 2006; Kruse and Julicher, 2000) flows are immediately expected in migrating cells. Retrograde flow of the F-actin network in the lamellipodia meets the anterograde flow of cell body F-actin in the so-called “convergence zone” (Salmon et al., 2002), resulting in anti-parallel F-actin movements and/or F-actin depolymerization. To investigate whether overlapping contractile F-actin flow patterns exist in migrating epithelial cells, we re-analyzed previously published FSM datasets of PtK1 cells (Gupton and Waterman-Storer, 2006). Previous analysis (Ponti et al., 2005) of F-actin dynamics identified the presence of two anti-parallel F-actin flow fields - a retrograde flow (Heath and Holifield, 1991) in the lamella and lamellipodia, and a forward F-actin movement in the cell body (Gupton and Waterman-Storer, 2006). Disappearance of flow vectors in the so-called “convergence zone”, where the two anti-parallel flow fields meet, was interpreted as F-actin disassembly, and previous tracking methods have not provided evidence of any anti-parallel (contractile) F-actin movement in the “convergence zone”. However, a visual inspection of these FSM image sequences shows that the method fails to correctly track the direction and speed of the overlapping F-actin flows (yellow vectors on the inset at the upper right corner of Fig. 3A).

**Figure 3.**
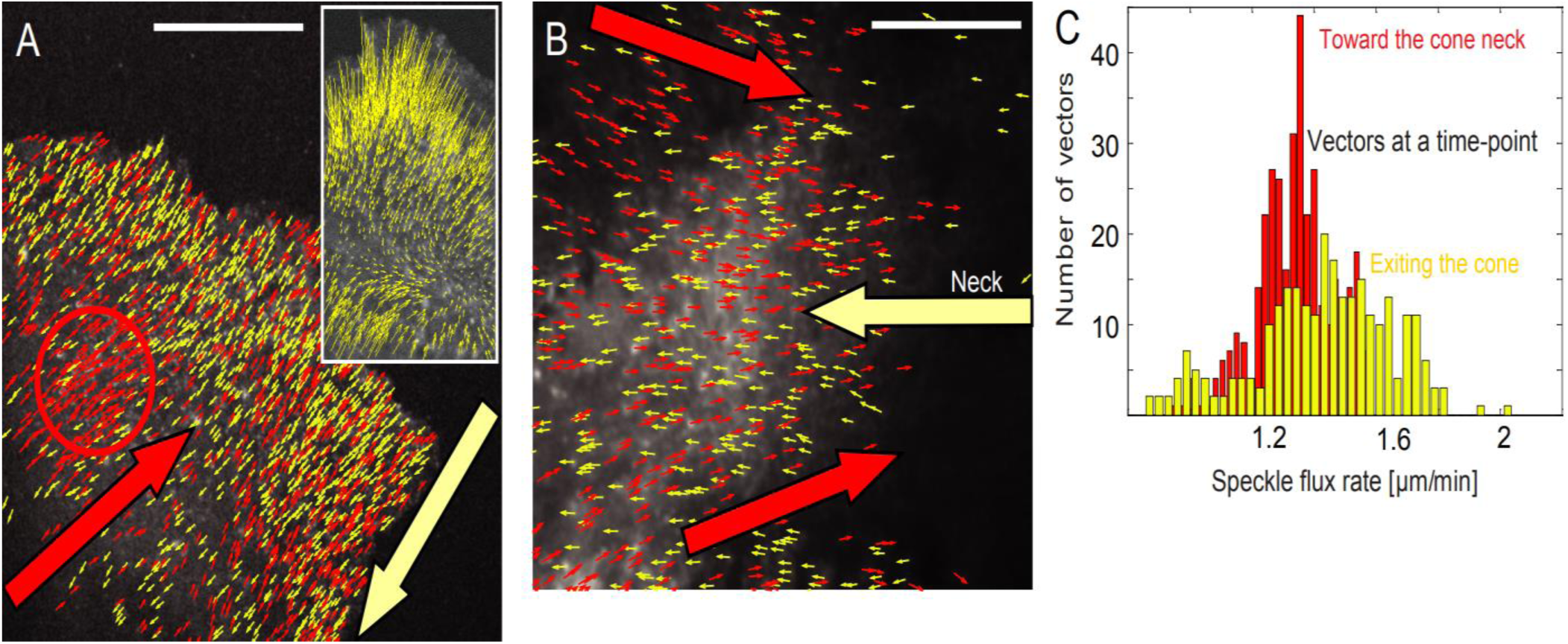
Actin networks with anti-parallel speckle flows. IFTA was used to track bi-directional F-actin polymer flux in migrating kangaroo rat kidney epithelial PtK1 cells and growth cones in primary cultures of *Aplysia bag* cell neurons imaged using FSM. Figure 3A. Contractile actin networks. We measured a considerable overlap of the two interdigating networks and, specifically, emerging anti-parallel flows in the intracellular regions previously thought to be zones of F-actin depolymerization (based on an earlier flow analysis presented in the inset of this figure). It is of significant difficulty to detect any zone of absolute convergence in this dataset, that is, an area of no speckle flux and diffusion of free dimers, as suggested by the measurement depicted by the speed vectors of length close to zero inset (see the yellow vectors showing no flux in the so-called “convergence zone”, based on an earlier analysis over the full image sequence) taken from (Gupton and Waterman-Storer, 2006). The tracking method used in (Gupton and Waterman-Storer, 2006) cannot handle anti-parallel motion and vectors with opposite directions, belonging to opposing flow fields, cancel out when directional filtering (that, spatial regularization) is performed during data postprocessing. This inaccuracy in the quantification has so far been interpreted as a local depolymerization activity. Using IFTA, we can now reliably measure that the depolymerization rates in the “convergence zone” are significantly lower than previously appreciated. Scale bar equals 5 µm. Figure 3B. Anti-parallel flux in the base of the growth cone. The flows in the two directions have different speeds and the retrograde flux is on average slower, but more homogenous (see the histograms in Fig. 3C). The image datasets were taken from (Burnette et al., 2007). In growth cones, two anti-parallel F-actin networks are thought to merge at the base, but visual inspection points at slight contractions (see also Vid. 6). Growth cones have a so-called “convergence zone” near the axonal shaft and previous analyses have not resolved the overlapped actin flows at the so-called neck of the cone. We found that the anterograde and retrograde flows interdigitate and flux with different speeds in the range of 0.8 to 2.1 µm/min. The anterograde flow passing through the neck of the growth cone was oriented straight toward the center of the leading edge (observe that the yellow vectors have a high level of angular orientation homogeneity) exhibit a high level of speed heterogeneity (yellow arrows in Fig. 3B and yellow bins in Fig. 3C). Conversely, the retrograde flow showed a narrow (and also slower on average) speed distribution, but a higher heterogeneity in terms of angular orientation (red vectors), as the F-actin translocation followed the outer shape of the cell (red arrows in Fig. 3B and red bins in Fig. 3C). Scale bar equals 10 µm.

In this context, IFTA measured a considerable overlap of the two interdigating networks and, specifically, emerging anti-parallel flows in intracellular regions previously thought to be zones of F-actin depolymerization (a side-by-side comparison is presented in Fig. 3A and the inset with yellow vectors at the upper right corner of the figure). In the area marked with a red circle, for instance - similarly to (Gupton and Waterman-Storer, 2006) - we measured a dense anterograde flow of about 2 µm/min in each direction. In the area thought to be convergent with no flux (Vallotton et al., 2004), however, we measured two overlapped networks, an anterograde flow (shown in red vectors) and a retrograde flow (shown in yellow vectors). It is, in fact, of significant difficulty to identify any zone of absolute convergence (Goins and Mullins, 2015) in this dataset, that is, an area of no speckle flux and diffusion of free dimers, as suggested by the measurement displayed by the speed vectors of length close to zero in the inset in Fig. 3A (the yellow vectors are showing no flux in the so-called “convergence zone”, based on an original analysis over the full image sequence), taken from (Gupton and Waterman-Storer, 2006). Thus, although F-actin depolymerization certainly occurs in the so-called “convergence zone”, all previously measured depolymerization rates are likely very largely overestimated (Vallotton et al., 2004). The reason for this error is that anti-parallel flow fields cancel each other out when regional vector filtering is performed (Ponti et al., 2005) during the iterative data processing in the previous tracking analysis (Gupton and Waterman-Storer, 2006; Vallotton et al., 2004).

The tracking method used in (Gupton and Waterman-Storer, 2006) cannot handle anti-parallel motion and vectors moving in opposite directions, that is, belonging to opposing flow fields, cancel out when directional filtering is performed during data postprocessing. This inaccuracy in quantification has so far been interpreted as local depolymerization activity, which is incorrect. Using IFTA, we can now reliably measure that depolymerization rates in the “convergence zone” are significantly lower than previously appreciated, while the polymerization rates are considerable. Conversely to the previous analyses, our measurements showed that the two, a retrograde and an anterograde, interdigitated speckle flow fields are extending way beyond the so-called “convergence zone”. Taken together, the measurement results with our method demonstrate that the two anti-parallel actin speckle flows are clearly interdigitating (see also (Matov, 2025b)) and fluxing in opposite directions in the areas of network overlap, thus suggesting these areas are significantly contractile and a “convergence zone” does not exist (“the chorus is the verse”).

#### Anti-parallel actin flows in the “convergence zone” at the base of neuronal growth cones

In neuronal growth cones, the F-actin network (Gallo et al., 2002) is organized similarly to that at the front of migrating cells and we next asked whether IFTA can dissect the arrangement of contractile F-actin networks in migrating neurons. In growth cones, two anti-parallel F-actin networks are thought to merge at the base (Burnette et al., 2007), but visual inspection of the imaging data indicates clear contractions at the narrow entry point at the neck of the growth cone (Vid. 6). In this context, a “convergence zone” was thought to exist near the axonal shaft, because previous analyses (Burnette et al., 2008) were not able to measure the velocities of the F-actin flows at the growth cone neck. Using IFTA, we were able to identify clearly present anterograde and retrograde flows, which were interdigitated (Matov, 2024h).

We measured heterogeneous flux rates - and direction of motion - with actin speckle speeds (Fig. 3B) in the range of 0.8 to 2.1 µm/min (Fig. 3C). The flows in the two directions exhibit different speeds and the retrograde flux is on average slower, but more homogenous (see the histograms in Fig. 3C). The anterograde flow passing through the neck of the growth cone was oriented straight toward the center of the leading edge (yellow vectors in Fig. 3B, pointing to the left, have a high level of angular orientation homogeneity) exhibit a high level of speed heterogeneity (yellow arrows in Fig. 3B and a distribution shown in yellow bins in Fig. 3C). Conversely, the retrograde flow showed narrow (and also slower on average) speed distribution with a higher heterogeneity in terms of angular orientation (red vectors in Fig. 3B, pointing to the right, have a high level of angular orientation heterogeneity), as the F-actin translocation followed the outer shape of the cell (red arrows in Fig. 3B and a distribution shown in red bins in Fig. 3C). See Table 1 in Materials and Methods for further information regarding the conditions and perturbations.

Both results have a very intuitive interpretation as, at the cone (see also examples in (Matov, 2025b)), the anterograde F-actin polymer network translocates from a narrow zone of high density of parallel speckle motion to a wider zone within the growth cone - hence the vectors are mostly parallel, yet speckles move at a high variety of different speeds. Conversely, the retrograde flow showed a narrower – and a slower on average - speed distribution (that is, the high speed speckles were suppressed) but with a high level of heterogeneity in terms of angular orientation, as the F-actin translocation followed the outer shape of the cell (Fig. 3B, red arrows pointing to the right with a distribution shown in red bins in Fig. 3C). These results are also very intuitive in terms of a geometrical interpretation as speckle flux rates are reduced at the narrow entrance of the neck of the cone and, at the same time representing a point of entry, the wider space becomes very narrow - hence the higher angular homogeneity of the F-actin network entering the neck.

Besides F-actin networks in growth cones in untreated, control cells, we analyzed cells after the addition of 70 μM blebbistatin. Blebbistatin (Straight et al., 2003) is a small molecule inhibitor which inhibits the activity of myosin II (Toth et al., 2003). After treatment, we measured slower F-actin turnover rates and a reduction of the anti-parallel contractile motion in perturbed cells (Matov, 2024h; Matov, 2025b). Again, these new measurements seemed very intuitive given the effects myosin II filaments exert on the actin organization (Janson et al., 1992; Martens and Radmacher, 2008). Taken together, these new results provided us with confidence that IFTA can truthfully detect shifts in speckle velocity - both in terms of changes in the speeds and spatial orientation of the motion - for a complex organization of the actin flows.

#### Contractile multi-directional actin flows in the “convergence zone” in the area between the lamella and lamellipodia of epithelial cells with overexpressed tropomyosin-2

Tropomyosins are long coiled-coil dimers that bind along F-actin filaments (Cooper, 2002). In muscle cells, tropomyosins modulate myosin binding and activity (Lees et al., 2011; Wang and Coluccio, 2010). In non-muscle cells, increased amounts of tropomyosin alter lamella and lamellipodia dynamics (Gupton et al., 2005), but the underlying molecular mechanisms, and to what extent different tropomyosin isoforms differentially affect F-actin dynamics, are not understood. Anti-parallel F-actin flows are myosin-mediated and tropomyosins regulate myosin-mediated anti-parallel F-actin contractions (Kad et al., 2005; Verkhovsky et al., 1999; Wang and Coluccio, 2010). Indeed, excess tropomyosin in migrating cells has previously been shown to render the F-actin network hyper-contractile (Gupton et al., 2005). Further, tropomyosin prevents binding of both cofilin and the Arp2/3 complex to actin filaments in the lamella (DesMarais et al., 2002) and limits the size of the lamellipodia networks (Iwasa and Mullins, 2007). We, therefore, used our novel tracking method to measure how different tropomyosin isoforms modulate F-actin dynamics.

Concretely, we overexpressed two tropomyosin isoforms, one from the α-gene (tropomyosin-2 (Tm2)) and one from the δ-gene (tropomyosin-4 (Tm4)) in PtK1 cells. Visually, we observed highly dynamic, re-organizing F-actin networks. Tm2 has been shown to inhibit the formation of new actin filaments and further, to inhibit the Arp2/3-mediated branch formation (Blanchoin et al., 2001), suggesting that when it is overexpressed, actin turnover will preferentially occur along the already existing polymer meshwork.

We first applied a tracking method used in (Gupton et al., 2005) for actin speckle flow measurements to successfully detect and decouple the effects of oppositely organized F-actin meshworks (Ponti et al., 2004). It failed to measure the speeds in the area of overlap and displayed speed vectors of, close to, zero (similar to the example shown on the inset in Fig. 3A), as described in section above. Also, in both cases of overexpression of the two tropomyosin isoforms texture-based tracking (Ji and Danuser, 2005) yielded no interdigitating speckle flows (see the insets with yellow vectors in Fig. 4A and Fig. 4C), but rather displayed vectors of a retrograde flow at the cell edge and a zone of no flux in the lamella, both of which were deemed incorrect after visual inspection of the imaging videos. We then used IFTA to quantify (Tables 2 and 3) the effects of tropomyosin overexpression on F-actin dynamics.

**Figure 4.**
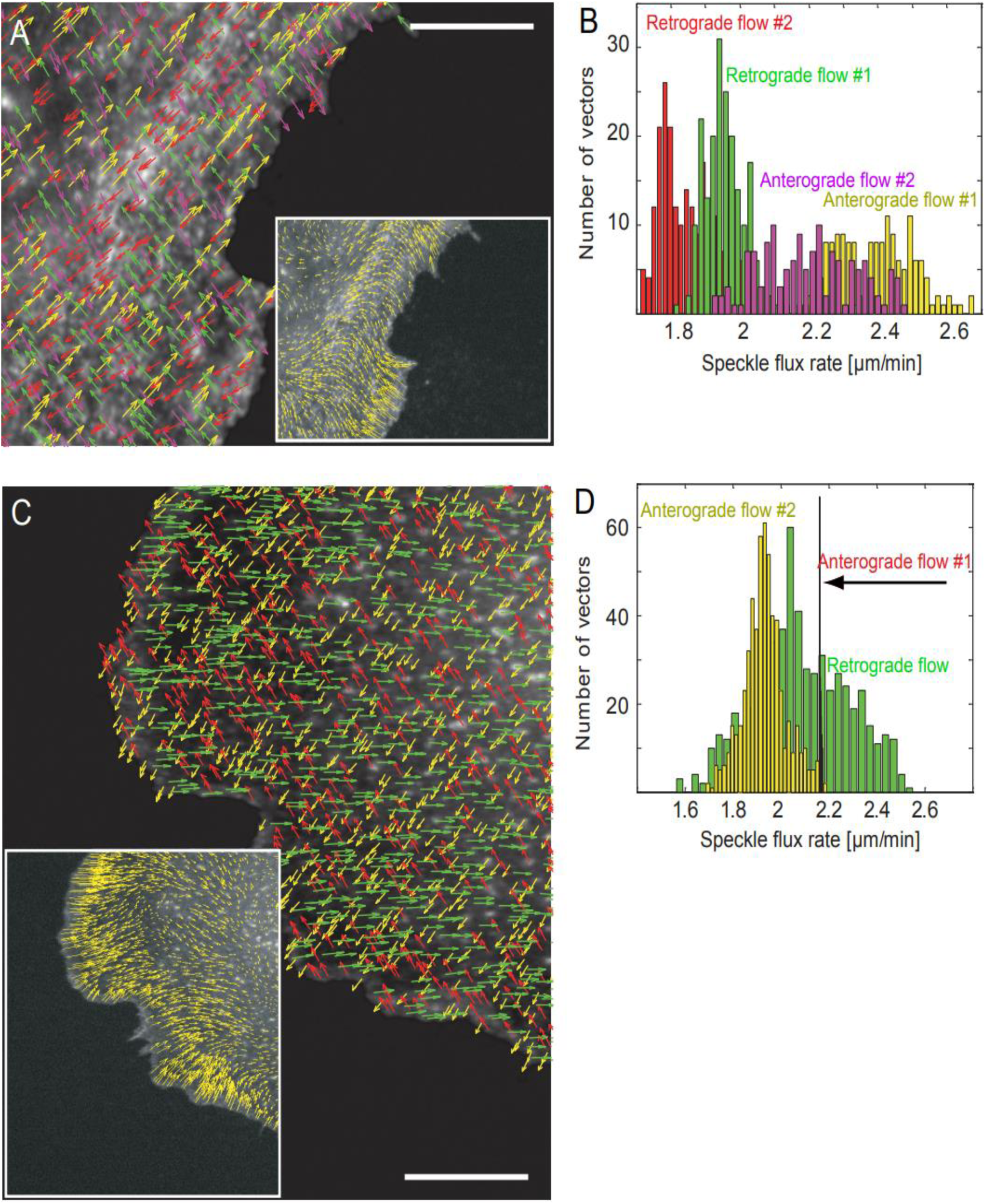
Highly contractile actin networks and overexpression of Tm2 and Tm4. IFTA was used to track four-directional and three-directional F-actin polymer flux in migrating kangaroo rat kidney epithelial PtK1 cells with overexpressed long (Tm2) and short (Tm4) human tropomyosin isoforms and imaged using FSM. Figure 4A. Tm2 overexpression increases anti-parallel left-right actin flux and leads also to anti-parallel up-down translocation, that is, a four-directional actin flow. We identified examples of four overlapping flows in cells overexpressing Tm2 (Figure 4B shows the corresponding four-color histogram) which seem to be coupled in pairs (see also Vid. 7, Vid. 8, Vid. 9, Vid. 10, and Vid. 11). Two of these flow fields exhibit close to perpendicular (forming an angle of 74°) to each other anterograde type of motion (in magenta and yellow vectors) with faster flow speeds and higher level of heterogeneity and standard deviation of the speed distribution (see the histograms in Fig. 4B). Conversely, the two retrograde flows (in red and green vectors, forming an angle of 73°) have slower and narrower distributions; those two flows (the one represented with red arrows in particular) seem to stem from an overall underlying polymer translocation rather than an active F-actin meshwork activity, as the speed distributions have lower standard deviation, the reason for which might be that the whole structure is moving and, thus, all associated speckles move with very similar speeds. This observation, together with the high edge activity, compared with wild type cells, suggests that Tm2 plays a role in the fast remodeling of F-actin filaments at the edge, yet preserves the shape of the speed distributions. Tm2, thus, works in this scenario like an F-actin filament stabilizer and long F-actin filaments decorated with Tm2 are stable and, therefore, can lead to the formation of protrusions in new, different directions. In comparison to IFTA, texture-based tracking (Ji and Danuser, 2005) yielded no interpenetrating speckle fluxes, but rather displayed vectors of a retrograde flow at the cell edge and a zone of no flux in the lamella, both of which are incorrect (see the inset with yellow vectors in Fig. 4A). Scale bar equals 5 µm. Figure 4C. Tm4 overexpression makes actin flux more heterogeneous (Vid. 12). In the case of Tm4 overexpression, actin speckle flows often exhibit two peaks in the distribution. The two “waves” of speckles resemble a distribution in which, first, there is a distribution similar to the wild type speeds and then, there is a second, faster peak centered around the average F-actin flux speed for data with overexpressed Tm2. Hence, many of the Tm4 overexpression datasets demonstrated an incorporation of both a wild type behavior and a Tm2 overexpression behavior. We also observed that while the retrograde or anterograde speeds of the F-actin flows were increased with the overexpression of Tm2, the overexpression of Tm4 created visibly different flow patterns with multi-directional flows where a sub-population behaves like wild type, while another one moves faster and in a distinctly heterogeneous way. Tm4 is a weaker actin binder and therefore its overexpression can stabilize nascent F-actin filaments. In the histograms shown in Fig. 4D (in three colors – green, red, and yellow), one can observe the very narrow peak formed by a flow moving parallel to the cell edge (red arrows and red bins). Remarkably, with values ranging 2.16-2.18 µm/min, there is a clear indication of a highly organized motion with an extremely narrow speed and angular variation, and standard variation. In comparison to IFTA, texture-based tracking (Ji and Danuser, 2005) yielded no interpenetrating speckle fluxes, but rather displayed vectors of a retrograde flow at the cell edge and a zone of no flux in the lamella, both of which are incorrect (see the inset with yellow vectors in Fig. 4B and Vid. 12). Scale bar equals 5 µm. Overall, in contrast to Tm2 overexpression, the retrograde flow (in green color in Fig. 4C and Fig. 4D) is faster than the anterograde, ranging from values lower than the anterograde to significantly faster. The speed vectors and histograms are based on the analysis of the image triplet consisting of time-frames 5-6-7 (20 seconds earlier than the analysis results shown in Table 2, which is based on time-frames 7-8-9).

**Table 2.**
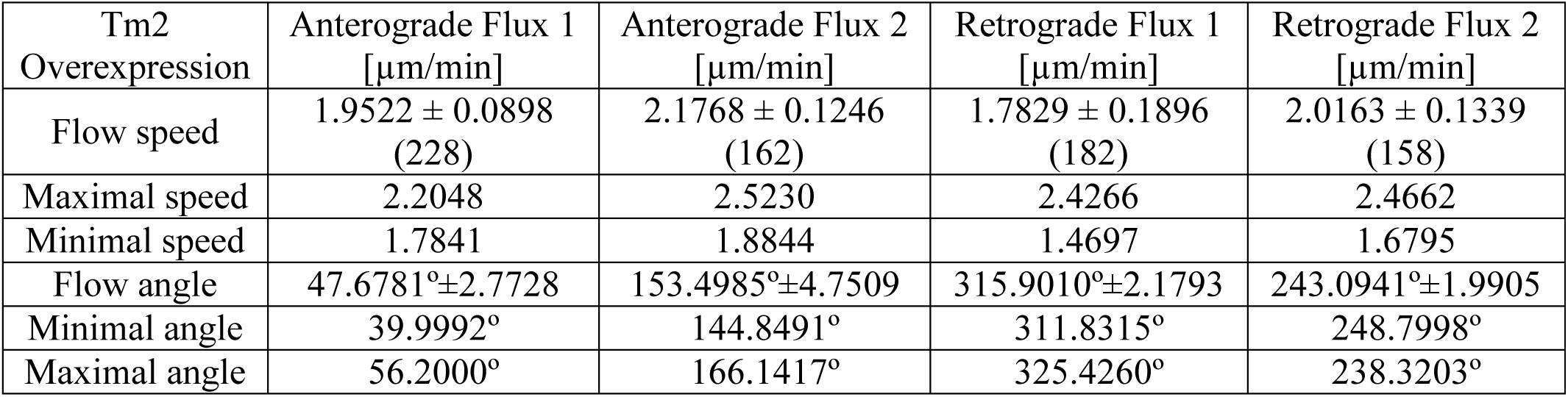
Four-directional flux in PtK1 cells overexpressing tropomyosin-2. See Vid. 7. Median flux speed values for the anterograde flow 1 (first column; see also the yellow vectors in Fig. 4A and Vid. 8), the anterograde flow 2 (second column; see also the magenta vectors in Fig. 4A and Vid. 9), the retrograde flow 1 (third column; see also the green vectors in Fig. 4A and Vid. 10), and the retrograde flow 2 (fourth column, see also the red vectors in Fig. 4A and Vid. 11) are presented together with a standard deviation. The numbers in brackets show the number of vectors tracked for one triplet of images (out of the 119 images in the time-lapse sequence) only. The speed and angle numbers are based on the analysis of the image triplet consisting of time-frames 7-8-9 (20 seconds later than the analysis results shown in Fig. 4, which is based on time-frames 5-6-7). The median direction of the flows with a standard deviation as well as the minimal and maximal angle of the flux vectors in each flow are listed in the bottom three rows. The y-axis on Fig. 4A (or a vertical line pointing upward) is the zero degree direction and the degrees increase in the clockwise direction.

**Table 3.** Three-directional flux in PtK1 cells overexpressing tropomyosin-4. Median flux speed values for the anterograde flow 1 (first column; see also the red vectors in Fig. 4C), the anterograde flow 2 (second column; see also the yellow vectors in Fig. 4C), and the retrograde flow (third column; see also the green vectors in Fig. 4C) are presented together with a standard deviation. The numbers in brackets show the number of vectors tracked for one triplet of images (out of 119 images in the time-lapse sequence) only. The speed and angle numbers are based on the analysis of the image triplet consisting of time-frames 23-24-25. Color-coding (blue) indicates a very low standard deviation of the flux rate and the direction of flux, and (red) high flux rates. The median direction of the flows with a standard deviation as well as the minimal and maximal angle of the flux vectors in each flow are listed in the bottom three rows. The y-axis in Fig. 4C (or a vertical line pointing upward) is the zero degree direction and the degrees increase in the clockwise direction.

| Tm4 Overexpression | Anterograde Flux 1 [μm/min] | Anterograde Flux 2 [μm/min] | Retrograde Flux [μm/min] |
| --- | --- | --- | --- |
| Flow speed | 2.1660 ± 0.0023 (462) | 1.9316 ± 0.0913 (581) | 2.0695 ± 0.1957 (570) |
| Maximal speed | 2.1712 | 2.1948 | 2.5539 |
| Minimal speed | 2.1633 | 1.6886 | 1.5619 |
| Flow angle | 336.6657° ± 0.0898 | 245.7843° ± 4.1746 | 82.4929° ± 2.4382 |
| Minimal angle | 336.6180° | 231.4589° | 75.8652° |
| Maximal angle | 336.9009° | 255.9540° | 89.9960° |

Our measurements indicated that Tm2 overexpression increases anti-parallel, left-right actin flux and leads also to anti-parallel, up-down translocation, that is, an overall four-directional actin flow. Using IFTA, we measured that Tm2 overexpression increases the speed of F-actin flux by about 30%, with speeds ranging from around 1.6 µm/min (speeds that are similar to those in wild type cells) to 2.7 µm/min, while preserving the organization of F-actin in retrograde and anterograde flows that interdigitate at the so-called “convergence zone”. We identified examples of four overlapping flows (Fig. 4A) in cells overexpressing Tm2 (in Fig. 4B, see the speeds of four vectors flows in the color histogram), which seemed to be coupled in anti-parallel pairs (see the tracking vectors for all four flows in Vid. 7 and individually for each of the four flows in Vid. 8, Vid. 9, Vid. 10, and Vid. 11). None of the four speckle flow fields was a result of a stage drift. We established that there are four significant flows by using expectation-maximization based automated selection of the number of vector cluster directions (Figueiredo and Jain, 2002; Matov et al., 2011), an algorithm that compares the iterative clustering of the vectors in one to nine flows and identifies the best match.

Two of the flows exhibit close to perpendicular (around 70°, Table 2) to each other anterograde type of motion (in magenta and yellow in Fig. 4A and Fig. 4B) with faster flow speeds and a higher level of heterogeneity and standard deviation of the speed distributions (see the histograms in Fig. 4B). Conversely, the two retrograde flows (in red and green vectors in Fig. 4A and Fig. 4B), similarly forming an angle of about 70° (Table 2) exhibit slower and narrower speed distributions, as the speed distributions have lower standard deviation, the reason for which might be that the whole actin meshwork is moving forward and, thus, all retrograde flow vectors seem to exhibit very similar flux rates (likely, slightly underestimated, also because of the overall motion in the opposite direction toward the cell edge). The organization of the two retrograde flows (the one represented with red arrows in Fig. 4A, in particular) seem to stem from an overall underlying polymer translocation rather than an active F-actin meshwork contraction. That retrograde flow speed distributions exhibit a lower standard deviation, the reason for which might be that the whole structure is moving and - with that polymer translocation - all speckles move with very similar speeds. This observation, together with the high edge activity compared to wild type, control cells, suggests that Tm2 plays an essential role in the rapid remodeling of F-actin filaments at the cell edge. Yet, its overexpression preserves the shape of the speckle speed distributions, that is, preserves the overall organization of the actin meshwork. Our new results indicate that Tm2 acts as a potent actin filament stabilizer and that the long actin filaments decorated with Tm2 are stable and, therefore, can lead to the formation of protrusions in multiple directions.

#### Contractile multi-directional actin flows in the “convergence zone” in the area between the lamella and lamellipodia of epithelial cells with overexpressed tropomyosin-4

We also studied the effects of overexpressing Tm4, another tropomyosin isoform. In contrast to Tm2 overexpression, in cells overexpressing Tm4, the retrograde flows are on overall faster than the anterograde flows. The retrograde speeds range from values lower than the anterograde to significantly faster speckle speeds of almost 2.6 µm/min, which are 3.6 µm/min faster than any anterograde moving speckles (Fig. 4 and Table 3). While the distributions of the retrograde and anterograde speeds of actin speckles in cells overexpressing Tm2 are normally distributed and exhibiting unimodal, normal distributions that are faster than in wild type data, a typical feature after the overexpression of Tm4 is the formation of actin speckle flows that exhibit two peaks and form bi-modal distributions. Further, in contrast to Tm2 overexpression, the retrograde flow in Tm4 overexpressing cells (in green color in Fig. 4C and Fig. 4D) is faster than the anterograde flows – however, it is very heterogeneous, with some speckle flux values lower than the anterograde flows to speeds that are about 20% faster. We can describe the retrograde flow speed distribution in Fig. 4D (in green) as having two “waves” of speckle vectors and resembling a distribution in which there is a first, slower mode - similar to the wild type speeds - and there is also a second, faster mode (or peak) centered around the average F-actin flux speed in some of the cells with overexpressed Tm2. Hence, many of the Tm4 overexpression image sequences demonstrated actin dynamics of embedded both wild type behavior and Tm2 overexpression behavior. See Table 1 in Materials and Methods for further information regarding the conditions and perturbations.

We observed that if the overexpression of Tm2 increased speeds of the retrograde or anterograde F-actin flows, the overall shape of the speed distributions remained similar to those in wild type cells. However, Tm4 overexpression created distinctly different flow patterns of three-directional flows, which consist of sub-populations behaving like in wild type cells and other sub-populations that move faster and in a distinctly heterogeneous way. Furthermore, in cells overexpressing Tm4, we often observed drastic directional changes in the flow organization within the 10 minutes duration of the acquired movies. It seemed that the zones of actin flow overlap were undergoing significant remodeling as a result of the induced contractions. Conversely, such changes never occurred in cells overexpressing Tm2 and in such cells the organization – in terms of the shape of the distributions of the speeds within the flows - remained unchanged in the zone of flow overlap.

Tm4 is the weaker and more transient actin binder (Gunning et al., 2005), and therefore its overexpression can stabilize nascent F-actin filaments, but it has lesser effects on the organization of the retrograde F-actin meshwork, that is, a smaller number of organized retrograde flows are visible, yet all of them exhibit an increased heterogeneity of the distribution of speckle flux speeds. Our results also indicate that Tm2 overexpression allows us to distinguish at a higher degree the branched, overlapped F-actin networks and that can result in the analysis of four overall actin flows – while in wild type cells are visible only two anti-parallel flows.

Remarkably, in the histogram shown in Fig. 4D (the three color speed distributions), one can observe a very narrow peak formed by a flow moving parallel to the cell edge (red upward vectors in Fig. 4C); with values ranging from 2.16 to 2.18 µm/min, a clear indication of a highly organized motion with an extremely narrow speed and angular variation (Table 3). In comparison to IFTA, texture-based tracking (Ji and Danuser, 2005) yields no interpenetrating speckle fluxes, but rather displayed vectors of a retrograde flow at the cell edge and a zone of no flux in the lamella (see the inset with yellow vectors in Fig. 4C), both of which are incorrect (Vid. 12). We did not observe any stage drift nor any movement of the cell body and therefore, this measurement was not a result of an artifact, but rather represents a new result demonstrating the ability of Tm4 to induce a very coordinated contraction parallel to the protruding cell edge.

Very similarly, protruding metastatic B16 melanoma cells form three actin networks in the lamellipodia, one of which consists of bundled F-actin in parallel to the cell edge (Koestler et al., 2008). The mechanism proposed consisted of alternating formation of two diagonal networks during protrusion and of parallel to the cell edge bundles during retraction (or withdrawal). Because of an improved visualization and analysis methodology, our analysis demonstrates that all three networks are active concurrently (Fig. 4C). Furthermore, our analysis demonstrates that the angle between the retrograde (73°) and anterograde (74°) networks is about 70° (Table 2), which is consistent with previously published data (Mullins et al., 1998; Svitkina and Borisy, 1999), when Tm2 is overexpressed. In the cell with overexpressed Tm4, the angle between the anterograde networks is about 91°. However, that perceived increase in the angle is due to the edge protrusion in upward direction (see the remarkably low standard deviations of both the flux speeds and the angles of the vectors of the first anterograde flow in Table 3). Further, the very high variability of the flux speeds distribution of the retrograde flow (the green distribution in Fig. 4C) suggests the presence of two distinct networks with somewhat similar flux rates, which appear indistinguishable (Vid. 12). Thus, we suspect that the (undetected) second branch of the retrograde network is the branch that is anti-parallel to the first anterograde flow (shown with red vectors in Fig. 4C).

Our results suggest the activity of two branched, overlapped, anti-parallel actin networks (for a mechanistic explanation, see (Matov, 2025b). The unique ability of IFTA to decouple the activity of overlapped, anti-parallel networks allows us to identify the different flows, while previous methods – in which the readout captures only the dominant flow at any given moment – report angles from 15° to 90° in the orientation of the filaments (Koestler et al., 2008), which we find unlikely, since the architecture of the actin meshwork is known (Mullins et al., 1998; Svitkina and Borisy, 1999). In our opinion, what changes rapidly is not the angle of the actin filaments, but rather the rates of polymerization in the different branches of the overlapped, anti-parallel networks. This way is created an illusion of shifts in the direction of motion of the fluorescent probes in homogeneous polymer labeling imaging data (Koestler et al., 2008), which can be overcome by the utilization of a low labeling ratio (∼0.25%) of actin subunits (Waterman-Storer et al., 1998). In the speckling image sequences, one can clearly observe the incredibly rapid changes in the flux rates and pinpoint areas of the cell in which otherwise, in data with homogeneous labeling, no directed motion can be detected. To provide a quantitative example, in Fig. 4B, the two anterograde flows exhibit more heterogeneous speed distributions and are about 0.5 µm/min faster than the two retrograde flows, suggesting cell protrusion. In Table 2, which shows measurements collected 20 seconds later, conversely, the two retrograde flows exhibit more heterogeneous speed distributions and the overall speed differences are reduced, suggesting a transition to edge retraction.

The analysis of Tm2/4 datasets demonstrated that IFTA could detect differential effects of tropomyosin isoforms on actin contractility. Firstly, we concluded that overexpression of the Tm2 isoform, which is a stronger binder, causes a stable organization of the F-actin flows, resulting in visible flows in multiple directions with varying speed distributions and directions of flux. Thus, our analysis approach allows us to investigate novel mechanisms regulating contraction. Drugs to improve heart muscle (Muthuchamy et al., 1993) and other muscle activity can be tested *ex vivo* based on this quantitative assay. Secondly, the overexpression of the Tm4 isoform, the weaker actin binder, which is a more transient binder, has lesser effects on the organization, that is, causes a smaller number of visible flows, yet the flows seemingly exhibit an increased heterogeneity of the distributions of speckle flux speeds. Drugs to improve bone repair and brain function can be tested *ex vivo* based on our quantitative assay. This type of measurements could serve as a quantitative readout in functional drug testing in the context of bone remodeling. Osteoclast activity is reduced with lowering the levels of Tm4, leading to the inhibition of bone resorption (Park et al., 2024). Putative anti-aging compounds could be tested *ex vivo* for their ability to maintain a heterogeneous actin flows orthogonal to the cell edge, while inducing coordinated, fast polymer translocation in parallel to the cell edge. Our quantitative assay could be used to evaluate compounds to treat and prevent diseases of the brain ranging from stroke to dementia (Pleines et al., 2017).

The ability of IFTA to detect subtle differential effects in the organization of F-actin meshworks in areas of network overlap demonstrates the power of our quantitative method. Overall, our analysis demonstrates that, besides the two distinct F-actin networks driving the protrusion at the leading edge (Ponti et al., 2004), in migrating cells there are also multiple, distinct contractile F-actin networks in the so-called “convergence zone”, that is, at the interface between the lamella and lamellipodia. Our new results demonstrate that IFTA is a valuable tool that can accurately resolve the organized dynamics of interconnected polymers of cytoskeletal proteins, such as tubulin and actin, in the context of drug development. Finally, our work indicates that there exists a high level of complexity in the mechanisms regulating the areas of the cell where the cell edge ends and the cell body begins, and strongly suggests the existence of putative drug targets.

To continue our work on speckle clustering, we can analyze the organized motion of a speckle cohort as a broadcast channel with memory and present it in terms of Gaussian multi-user parallel broadcast channels with identical code words (Matov, 1999; Tong et al., 2024). Considering the Markov chain of speckle formation as a degraded broadcast channel, considerations regarding the capacity region (Han, 1981) and its estimation (Bergmans, 1973) for a cohort of speckles could further aid the classification of a new speckle to a particular flow field.

Furthermore, in some of the examples considered, such as the MTs within the mitotic spindle, the speckle motion can be defined as non-Markovian, that is, the next positions in the trajectories depend not only on the present state, but on the past states as well. In this context, complete speckle trajectories can be computed using a Markovian approximation (Lindblad, 1976; Manzano, 2020). However, it is fair to assume that there is memory in the trajectories, which means that a Markovian assumption might not be applicable. This is due to strong environment couplings, correlations, and entanglement in the motion. A powerful tool for dealing with such systems is provided by the Nakajima-Zwanzig equation (Nakajima, 1958; Zwanzig, 1960). This projection technique has manifested its efficiency at the construction of the generalized master equation and investigation of non-Markovian dynamics.

### Patient-derived organoids

To translate our findings from cell line models to systems with a high level of clinical significance, we began culturing patient-derived organoids. The organoid culture system allows us to draw conclusions regarding patient treatment options based on the analysis of patient-derived cells *ex vivo*. Initially, we cultured metastatic castrate-resistant PC organoids derived from retroperitoneal lymph node metastasis with prior treatment with androgen deprivation therapy, bicalutamide, docetaxel, and carboplatin (Gao et al., 2014). We plated organoids with diameter 40-80 µm in 30 µL Matrigel drops and cultured them for five weeks, during which some organoids exceeded ½ mm in diameter (Fig. 5). The organoids had a characteristic round shape and were rapidly proliferating, and overall easy to culture. Since these drug-resistant organoids exhibited a mesenchymal gene expression signature, we tested therapeutic approaches to inhibit the activity of GPX4 (Hangauer et al., 2017) and demonstrated that small molecules can electively induce ferroptosis in tumor cells with acquired non-mutational drug resistance (Matov, 2025a).

**Figure 5.**
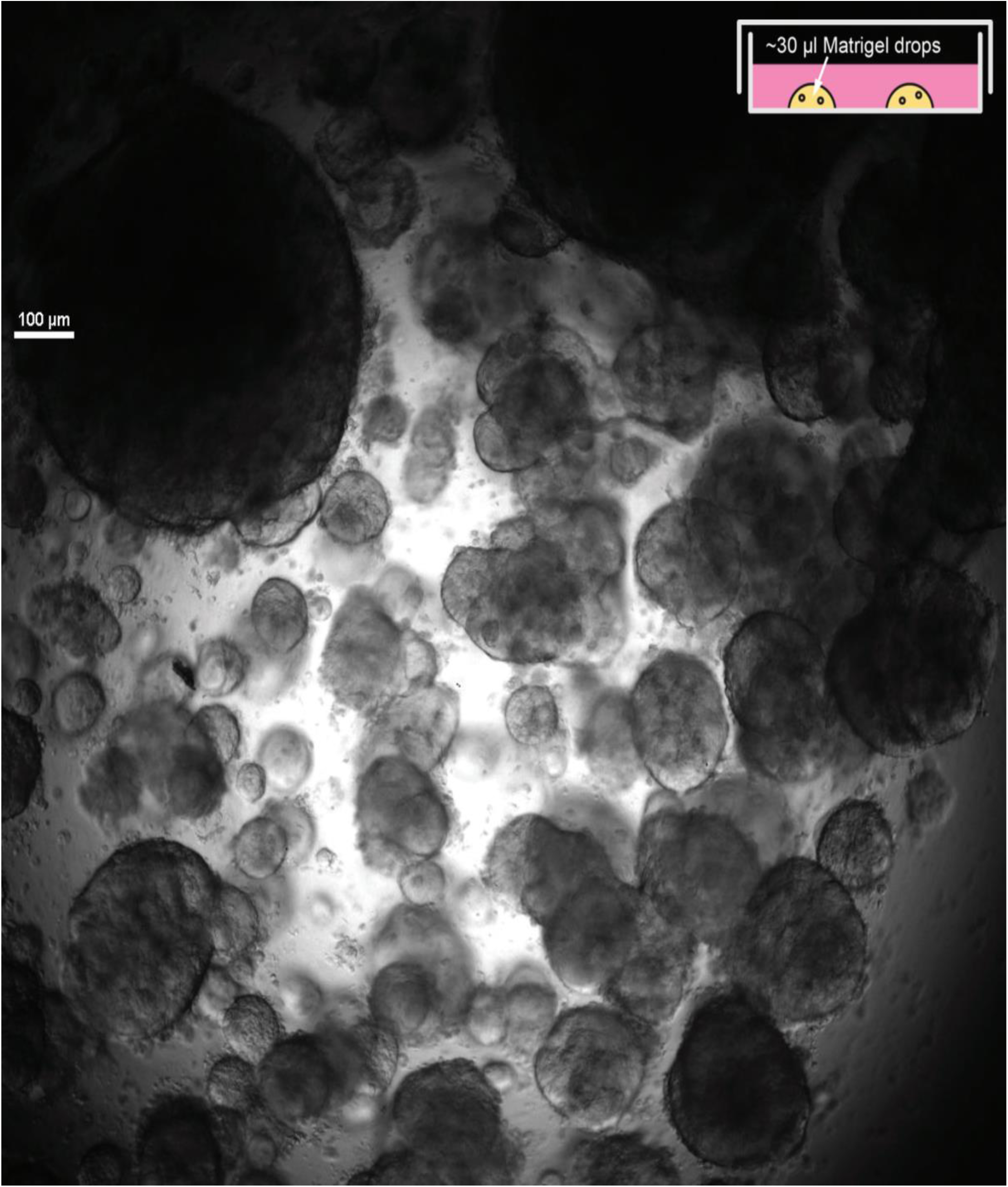
Metastatic prostate cancer organoids derived from retroperitoneal lymph node metastasis. Prior treatment – androgen deprivation therapy, bicalutamide, docetaxel, and carboplatin (Gao et al., 2014). Organoids with diameter 40-80 µm were plated in 30 µL Matrigel drops and cultured for five weeks. Transmitted light microscopy, magnification 4x. Scale bar equals 100 µm.

We derived organoids from high-grade acinar adenocarcinoma tumors. Our investigation demonstrated (Matov, 2024d) that high-grade primary prostate tumors are not reliant (Green, 2024) on the activity of the Rho-associated protein kinase (ROCK) to form organoids and we used ROCK inhibitors to select against prostate epithelial cells in our organoids culture. ROCK regulates myosin light chain phosphorylation (Chrzanowska-Wodnicka and Burridge, 1996; Katoh et al., 2001) and, as a result, actin activity and cellular contractility. While the use of ROCK inhibitors (Narumiya et al., 2000) impeded the ability of prostate epithelial cells to form organoids, the cells from high-grade primary tumors formed visible glandular structures, resembling two urethral sinuses, and acini (Fig. 6, Fig. S2, and Fig. S3). ROCK inhibitors inhibit the activity of myosin II, which reduces the contractility of the actin cytoskeleton (Matov, 2024h). GSK3α is elevated in low Gleason Score tumors, while GSK3β is overexpressed in high Gleason Score tumors (Duda et al., 2020), and the isoform-specific differentiation suggests that ROCK inhibitors can be utilized to select against early stage disease and prostate epithelial cells in organoid culture. GSK3β is downstream of RhoA (Narumiya et al., 1997) / ROCK and its overexpression in high-grade prostate tumors reduces the ability of ROCK inhibitors to affect its function.

**Figure 6.**
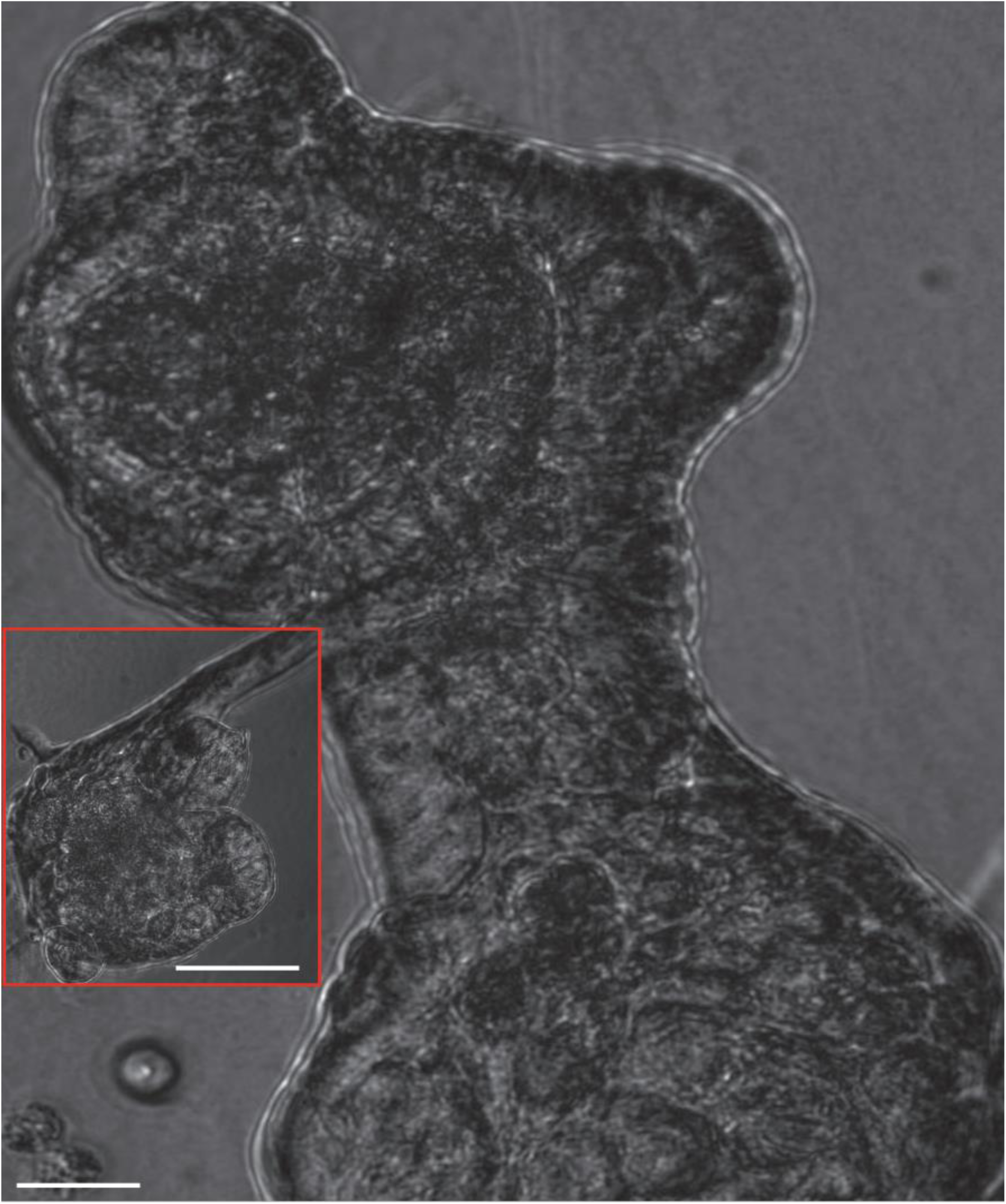
Acinar adenocarcinoma organoids obtained from organ-confined disease. Organoids derived from radical prostatectomy tissue. The organoids formed visible glandular structures, resembling two urethral sinuses. Organoid cells formed acini. Phase contrast microscopy, 20x magnification. Scale bar equals 10 µm. Inset scale bar equals 60 µm.

In our organoid culture, a concentration of 2.5 µM Y27632 (Darenfed et al., 2007) impeded the ability of prostate epithelial cells to form cell-cell junctions and form organoids without affecting the cells from primary prostate tumors, likely because of the overexpression of GSK3β in high-grade PC. Further, E-cadherin is overexpressed in high-grade prostate tumors (Matov, 2024d). FA-dependent acto-myosin traction force (that is, contraction) and tension on the cell–cell junctions disrupt cell–cell adhesion during epithelial cell scattering, which correlates with stronger FAs and increased phosphorylation of the myosin regulatory light chain (de Rooij et al., 2005). Further, stabilization of MTs in the vicinity of FAs is regulated by GSK3β inactivation and prostate epithelial cells cannot achieve cell polarity and migrate (Kumar et al., 2009) upon ROCK inhibition.

To measure the changes in PC cell spreading and cell edge shape, we segmented the cellular outlines of cancer cells with fluorescently labeled FAs after stimulation by HGF and an Epac (Grandoch et al., 2009) selective cAMP analog (Enserink et al., 2002), which activate the Ras GTPase Rap1 (de Rooij et al., 1998; Matov, 2024h). The induction of cell spreading in PC cells, decreased FA size and brightness, and inhibited FA sliding (Spanjaard et al., 2015) – similarly to the effects of inhibition of cytoskeletal contractility by RhoA, a downstream target of Rap1, a small GTPase very similar to Ras (Bos et al., 1997). Rap1 was observed to suppress oncogenic Ras phenotype and the proposed reason had been competition between Ras and Rap1 for a common, unidentified target (Kitayama et al., 1989). Rap1 is often mutated in PC (Matov, 2024g) and can promote metastasis (Bailey et al., 2009) as well as Rho signaling to recruit myosin to the cell edge (Rothenberg et al., 2023).

Breast epithelial MCF-10A cells exhibit faster MT polymerization compared to breast cancer cells (Matov, 2024d). Ras-transformed MCF-10A cells have elevated levels of myosin light chain phosphorylation and are more contractile than their normal counterparts, consistent with the activation of Rho. Furthermore, inhibitors of contractility restore an epithelial phenotype to the Ras-transformed MCF-10A cells (Zhong et al., 1997). The inhibition of contractility leads to the disruption of E-cadherin cytoskeletal links in cell-cell junctions and blocks the assembly of new cell-cell junctions. Some, but not all, characteristics of Ras-transformed epithelial cells are due to activated Rho. Whereas Rho is needed for the assembly of cell-cell junctions, high levels of activated Rho in Ras-transformed cells contribute to their altered cytoskeletal organization. Factors lowering the overall GTPase rate may increase the occurrence of GTP-tubulin caps at MT ends and GTP remnants along the MT lattice (Thoma et al., 2010), thus reducing the chances of improper chromosome segregation and aneuploidy. Several MT-interacting complexes are involved in the assembly of the mitotic spindle and regulated by RanGTP (Chang et al., 2009; Geraghty et al., 2021; Petry et al., 2013; Ribbeck et al., 2006; Silljé et al., 2006; Walczak et al., 1997) – NuMA, KifC1, TPX2, HURP, NuSAP, and astrin-kinastrin. RanGAP1 induces GTPase activity of Ran but not Ras, while the constitutively active mutant RanQ69L, which is unable to hydrolyze bound GTP as it is insensitive to RanGAP1, is locked in the active state (Bischoff et al., 1994). RanBP2 and RanGAP1 are involved in the disassembly of nuclear export complexes in the cytoplasm (Ritterhoff et al., 2016), while a pro-oncogenic noncanonical activity of a Ras-GTP:RanGAP1 complex has been shown to facilitate nuclear protein export (Tripathi et al., 2024).

We derived organoids from drug-naïve micro-metastatic tumor tissues resected from the retroperitoneal lymph nodes. Our observation has been that the required to recapitulate the tumor microenvironment cancer-associated fibroblasts (CAFs) - which we cultured with organoids from prostate tumors, since the fibroblasts can secrete tumor-specific growth factors (Cunha et al., 2002; Cunha et al., 2003; Cunha et al., 2004; Hayward et al., 1998; Olumi et al., 1999; Thomson and Cunha, 1999) - are not continuously activated in Matrigel. In our organoid culture, fibroblast cells (see examples from primary tumors in Fig. S4 and micro-metastatic tumors in Fig. S5) lasted for about two months and after that, only epithelial organoids remained (Fig. 7). Taxane treatment attenuates fibroblast motility in HMF3S cells and reduces the levels of lysyl oxidase homologue 2, c-Met, fibronectin (Huet-Calderwood et al., 2023), TGFβ, CTGF, importin α, and 14-3-3ζ in receptor triple negative breast cancer cells (Tran et al., 2013; Tran et al., 2012), suggesting approaches for modulating fibroblast activity in drug-treated tumors.

**Figure 7.**
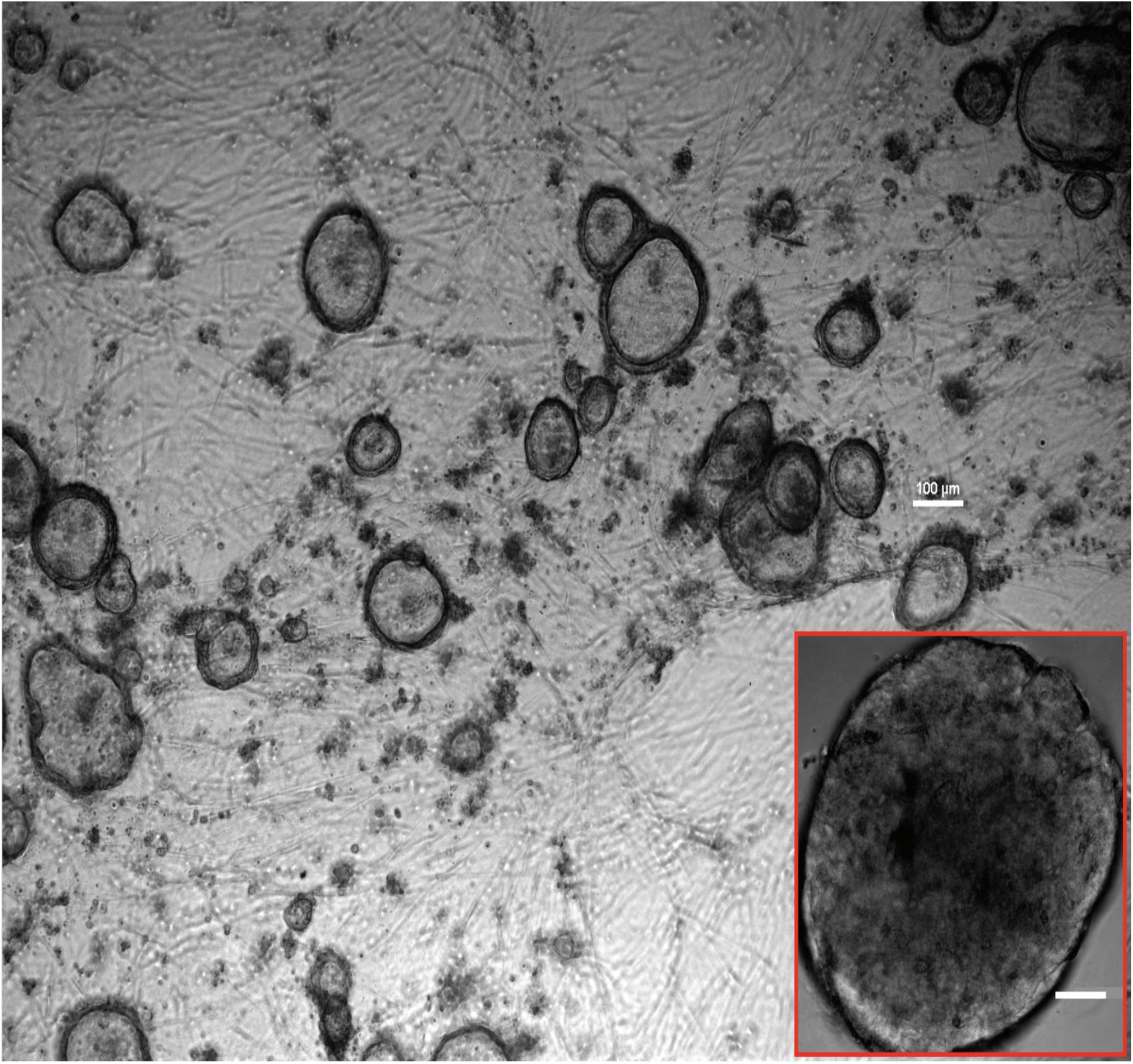
Metastatic prostate cancer organoids derived from retroperitoneal lymph node metastasis. Organoids derived from radical prostatectomy tissue. Drug-naïve tumor. Organoid whole genome sequencing showed homozygous mutations in *PTEN* and *BRCA2*. Day 84. Transmitted light microscopy, 4x magnification. Scale bar equals 100 µm. Inset: Day 73. Transmitted light microscopy, 20x magnification. Scale bar equals 50 µm.

The cofilin pathway is dysregulated in cancer and most commonly associated with the development of metastasis (Wang et al., 2007). Cofilins (Moon and Drubin, 1995) modulate actin (Welch and Mullins, 2002) dynamics at the leading edge of motile cells. Actin dynamics in lamellipodia are driven by continuous cycles of actin polymerization, retrograde flow, and depolymerization (DesMarais et al., 2005; Welch et al., 1997). Cofilin competes with tropomyosin to bind actin filaments and its activity is regulated by 14-3-3ζ (Gohla and Bokoch, 2002). 14-3-3ζ inhibits the dephosphorylation of cofilin, leading to increased tropomyosin binding along actin filaments, and suppresses actin turnover (Blanchoin et al., 2001). Cofilin activity directs TGFβ–mediated actin severing and stimulates cell migration (Bakin et al., 2004). The reduction of TGFβ and 14-3-3ζ after taxane treatment we measured would, thus, be associated with an increase in cofilin activity and a decrease in tropomyosin binding to F-actin, resulting in decreased ability of the cell for locomotion (Abercrombie et al., 1970).

The regulation of the prostate epithelial cells is mediated by the androgen receptor (AR). However, many “androgenic effects” on prostatic epithelium do not require epithelial AR, but instead are elicited by the paracrine action of AR-positive mesenchyme (Cunha, 2008). During prostatic tumorigenesis the stroma undergoes progressive loss of smooth muscle with the appearance of CAFs (Cunha et al., 2002). Paracrine stimulation causes an increase in epithelial AR and leads to tumorigenesis (Memarzadeh et al., 2007). Histologically, PC is defined by basal cell loss and malignant luminal cell expansion. Nevertheless, a paracrine growth factor can initiate tumorigenesis in basal cells as well (Lawson et al., 2010). Epithelial AR can function either as a tumor promoter or a tumor suppressor and its function in prostate tumorigenesis depends on the cancer initiating signal (Memarzadeh et al., 2011). This is particularly important in culturing prostate tumors as organoids. In our experience, paracrine signaling is paramount to organoid growth. Cultures that lose the CAFs, seize growth and become quiescent (Fig. 7).

In our organoid culture, the activation of CAFs took about 72 hours every time we refreshed the Matrigel. Such behavior indicates that soft Matrigel is an obstacle for mesenchymal cells and suggests co-culture of epithelial tumor organoids in suspension with the CAFs cultured below them without Matrigel. In this context, analysis of FSM image datasets with IFTA in organoids will allow us to measure changes in contractility and fine-tune the culture conditions.

The tumor microenvironment (Hanahan and Weinberg, 2011) exists as a 3D niche in which tumor cells lay on the extracellular matrix (Yamaguchi et al., 2005) together with CAFs (Kalluri and Zeisberg, 2006; Olumi et al., 1999), endothelial cells (Denekamp, 1982), and immune cells (Binnewies et al., 2018), and we cultured patient-derived organoid cultures in which, even if temporarily, all these components were present (Fig. S6, Fig. S7, and Fig. S8). Mechanical cues, such as the extracellular matrix stiffness, play a role in fibroblast activation and survival.

We performed lentiviral transduction with MT markers immediately after tumor tissue dissociation, before we seeded single cells in Matrigel, and the organoids expressed EB1ΔC-2xEGFP a day after radical prostatectomy. Our analysis of MT tip labeling with ClusterTrack indicated that the CAFs had 2-fold larger area and over 3.8-fold higher density of polymerizing MTs compared to the prostate tumor cells (Matov, 2024a). The high level of MT polymerization activity in CAFs (Vid. 13) and their extensive length (>100 µm) allow for a rapid paracrine signalling in prostate tumors. Our work has shown the extraordinary ability of small organoids to move in Matrigel and form large structures (Matov, 2025b) in samples from micro-metastatic tumors.

We derived organoids from castrate-resistant PC that had metastaized to the liver and in these cultures we also observed rapid clumping of tumor cells during the week after biopsy and organoid seeding (Fig. 8). What we noticed on the third day after seeding was that many of the small organoids had formed a large structure of 0.6 mm in diameter (Matov, 2024a; Matov, 2025b), which was remarkable. It underscores the incredible ability of these tumors to establish signaling communication in a new environment and protrude to enhance their microenvironment (Matov, 2024a) - all of which required about 72 hours. After the formation of a few large structures on Day 3, the remaining organoids exhibited significantly reduced proliferation rates, with only a small fraction (fewer than a quarter) of the organoids increasing in size. By Day 71, the median organoid size had reached 80 µm in diameter, which suggests that there were about 60 cells within the organoids. The large structures had disappeared, likely because the CAFs had died.

**Figure 8.**
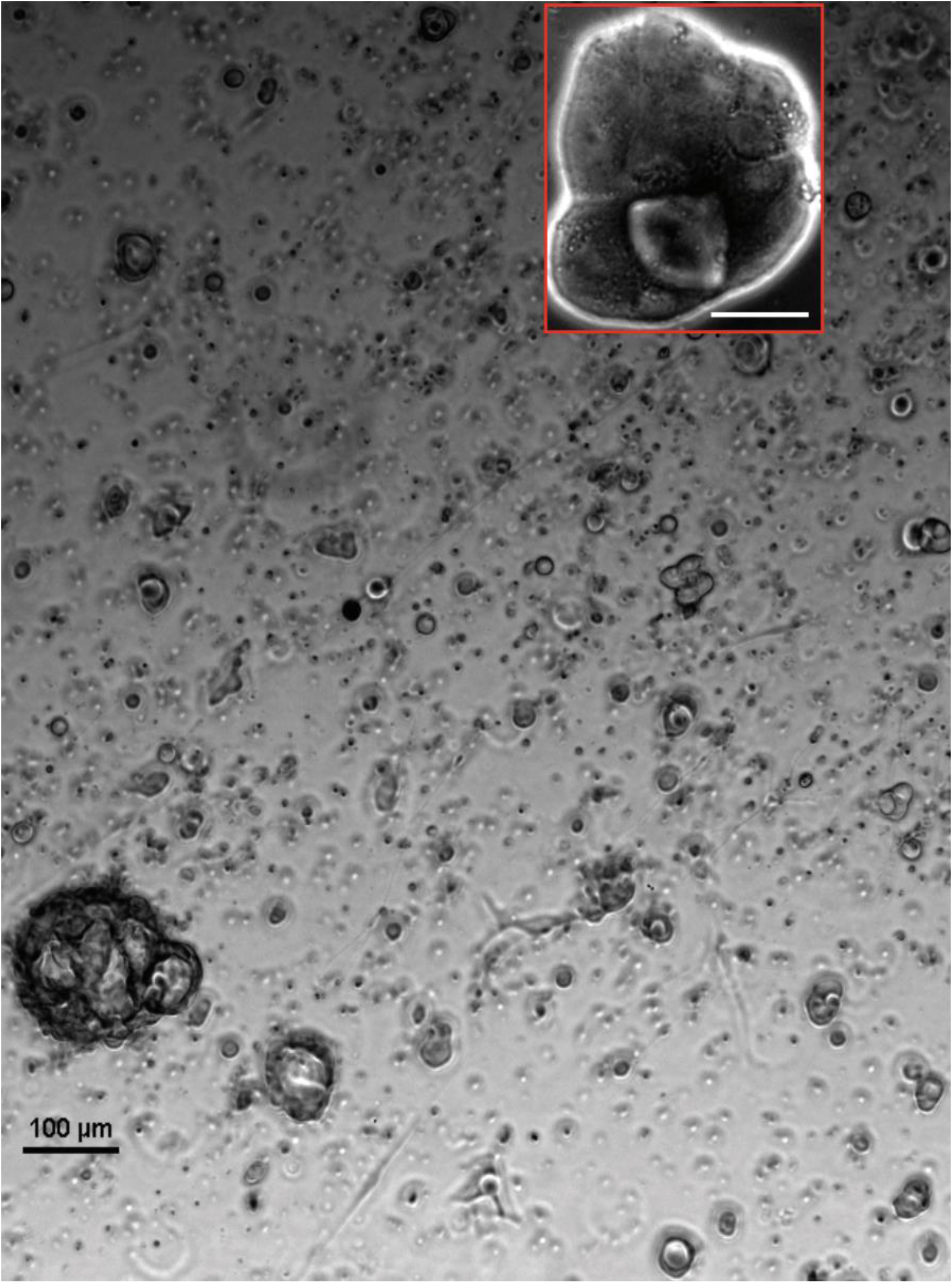
Metastatic prostate cancer organoids derived from liver metastasis. Organoids were derived from ½ mg needle biopsy tissue. Day 8. Transmitted light microscopy, 4x magnification. Scale bar equals 100 µm. The inset shows an organoid with irregular shape. Day 7. Phase contrast microscopy, 20x magnification. Scale bar equals 40 µm.

Fibroblast activation and deactivation affects the progression of PC (Owen et al., 2023) and the susceptibility (Smith et al., 2023) of prostate tumors to drug treatment. Contractility of activated fibroblast is reduced when myosin II is inhibited (Martens and Radmacher, 2008), which is the mechanism behind the partial - since there is a trade-off between the induced level of inhibition of cellular contractility and tumor prolifiration - success (Barcelo et al., 2023) of ROCK inhibitors in clinical trials. To calibrate the required drug combination, which deactivates CAFs but does not induce an increased resistance to apoptosis in the tumor, it would require analyzing the sub-cellular behavior in patient-derived cells. IFTA, the software approach we describe in this contribution, can aid the clinical determination of drug efficacy, as it relates to the contractility of key cytoskeletal components, such as the MTs and F-actin.

It is likely that misconceptions regarding the effects of the inhibition of Eg5/KIF11 are the result of wrong measurements (Miyamoto et al., 2004) - even if the contraction, in most cases, is clearly visible and has been pointed out by our collaborators and scientists who aquired the images we have re-analyzed. Similarly, the claims of an actin “convergence zone” at the interface between the cell body and the lamella as well as between the lamella and lamellipodia are puzzling. We have had discussions with a plethora of experimentalists who point at examples from their imaging datasets, which clearly indicate anti-parallel polymer flux, either due to MT or F-actin polymerization or polymer translocation activities. For these reasons, we have made an attempt to offer to the community of experimentalists our computational approach to resolving the complex and entangled motion at places in the cell where two distinct areas intersect. Our analysis can contribute to improving our understanding of cellular regulation not only in the context of fundamental research, but also by enhancing our ability to make clinical decisions pertaining to patient treatment in oncology and neurodegeneration.

## DISCUSSION

Until now, the accurate computational analysis of contractile cytoskeletal structures was limited due to the lack of appropriate software tools. Often speckle tracking in FSM images has been limited to the correct extraction of flow information at the cell edge up to the region of overlap of anti-parallel speckle flows. Since the entangled networks at the cell edge and in the cell body are fast evolving and rapidly changing their turnover rates and directions of motion, it has been impossible for traditional tracking methods to resolve the overlapped, anti-parallel motion. IFTA applies self-adaptive local and global motion models and is, hence, technically well equipped to analyze complex flow fields. Our method is very robust against frequent appearance and disappearance of speckles, as well as temporary occlusions, both characteristic of the dynamics in large contractile polymer structures with a significant turnover. This type of cellular flux measurements adds a new level of insight into the contractile cellular processes and advances our understanding of the dynamic organization of the cytoskeleton, cell division and motility, and thus, advances the field of drug development.

We have demonstrated the utility of IFTA for the resolution of multiple interdigitating flow fields of fluorescent fiduciary markers, termed speckles (Fig. 9). The goal of this contribution is to present examples of the analysis of cytoskeletal dynamics in which our approach resolves polymer flux that was previously missed by the global nearest neighbor (Ponti et al., 2005), texture-based flow tracking (Ji and Danuser, 2005), and projection-based particle tracking methods (Yang et al., 2005). The nearest neighbor method cannot handle high speed contractility measurements in high density networks. Such approach will always prefer to link speckles across oppositely oriented polymer fibers because of the high polymer density, thus erroneously resulting in short motion vectors (that is, close to zero speeds) and underestimated speckle flux rates (Vallotton et al., 2004) - and underestimated polymerization activity. The texture-based methods cannot cope with the task of tracking anti-parallel flows because of the utilization of an averaging step over multiple speckles in a region of interest, which leads to underestimating speckle speeds. The latter approach fails because of the latency of the linear Kalman filter (Swerling, 1959) applied to propagate the motion of speckles between consecutive frames. Thus, rapid flux changes either in space (change in direction) or time (change in speed, such as acceleration or deceleration) are often missed. We have, therefore, previously used IFTA to constrain the Kalman filter with information of spatio-temporal gradients in the instantaneous flow behaviors, which increased the Kalman tracking success rate from 61% to 96% (Matov, 2024i; Yang et al., 2005).

**Figure 9.**
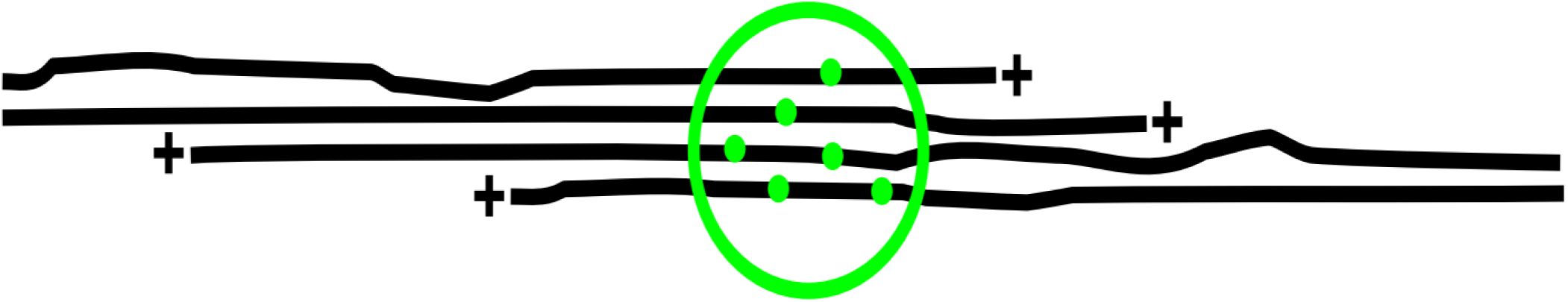
Speckle formation on MTs. In FSM images, when 4 to 7 fluorescently labeled tubulin dimers (green dots) fall within the diffraction limit of the microscope (about 220 nm in spinning disc confocal microscopy images), they form a bright speckle visible against image background and noise (Waterman-Storer et al., 1998); see the green circle on the figure. MTs are depicted in black and their polymerizing ends are marked with a (+) sign.

Our hypothesis that there exist overlaps – like Lego building blocks – between the actin networks in adjacent cellular areas seems very intuitive. According to feedback from our collaborators working on actin contractility, there is a clear polymer flux both in the areas of between the cell body and the lamella, and between the lamella and the lamellipodium. All results presented in this contribution are consistent with visual observations made by the experimentalists who acquired the imaging datasets. It is, therefore, difficult to conceive the reasons for the “convergence zone” hypothesis that has persisted in the field. It is true that some earlier computational analyses reported no flux at these “convergence zones”, but it is nevertheless clear that a visual inspection of the time-lapses sequences convincingly shows otherwise. It is also hard to imagine a complete seizure of polymer turnover and flux in living cells - such claims are simply contradicting reality.

Even in cells in which contractility is inhibited, we measured differential flux mechanisms. For instance, our measurements indicated that treatment with blebbistatin significantly reduced the anterograde flux in the “convergence zone” at the base of neuronal growth cones, without affecting the retrograde flux (Matov, 2025b). This new result suggests a gradient in myosin activity and our hypothesis is that myosin II expression is higher in the cell body compared to the area of the lamella (and its expression might be even lower in the lamellipodia (Vallotton and Small, 2009), creating an additional gradient in myosin activity). Further, analysis of vinculin speckles (Adams et al., 2004) after drug treatment can provide an additional quantitative readout regarding differential effects on the activity of FAs in the context of modulating the mechanisms of cell protrusion and retraction. Our new results of a differential effect and the preferential targeting of the actin anterograde network can be exploited in clinical applications in the field of neurology to stimulate axonal retraction and consequent neuronal regeneration.

In this context, IFTA adds a new level of insight, in addition to the already available information about speckle flows measured by earlier tracking methods (Ji and Danuser, 2005; Ponti et al., 2005; Yang et al., 2005). Undoubtedly, those three methods performed accurately and robustly the speckle tracking tasks they were designed for. IFTA, however, brings an additional strength as an add-on to those algorithms because of its capability to dissect specifically the cellular areas of active contraction, a key feature for many important cellular activities in disease. A technical shortcoming of IFTA, however, is that the algorithm extracts instantaneous flow behaviors, but does not assemble full speckle trajectories. Hence, the method is not suited for measurements when speckle lifetimes, from appearing or formation (“birth”) to disappearing (“death”), are required (see an example of such analysis in contact-inhibited cells in (Ponti et al., 2003)). However, in the scenarios we presented here, because of the rapid dynamics of the cellular polymers, the speckling fiduciary markers are only present for three to four movie frames (10 to 20 seconds). For this reason, IFTA is well adept to extract all of the available information in the image datasets presented in this contribution.

Importantly, our approach allows us to perform secondary, functional cellular assays and investigate the efficacy of the hits identified in primary high-throughput screening (Fox et al., 2006) assays in drug discovery. Such cell-based assays are designed for high-content screening (Nichols, 2007) of lead candidates, but the current image analysis methods have delivered limited precision. The utilization of IFTA in this context can drastically improve the quality of drug evaluation and increase the success rates of the clinical trials.

Measurements of cells overexpressing Tm2 and Tm4 demonstrate, in the new image datasets acquired for this manuscript, antagonistic effects on cellular contractility. The striking differences in the regulation of anterograde and retrograde F-actin networks, depending on the overexpression of the long or short isoform of tropomyosin, indicate potential applications of tropomyosin modulating medications in the regulation of heart function. Based on our analysis, Tm2 overexpression leads to the formation of two diagonal retrograde flows and two diagonal anterograde flows that are pair-wise anti-parallel. Further, Tm4 overexpression leads to the formation of three actin networks, one of which exhibits a retrograde flow with a wide variation in the flux speeds. We also measured completely opposite effects in regard to the speed of F-actin flux in retrograde (faster when Tm4 is overexpressed) and anterograde (faster when Tm2 is overexpressed) flows. Therefore, it is conceivable that these antagonistic effects may allow for a precise tuning of muscle contraction beyond maintaining cardiac health, but also to improve athletic performance and recovery.

Personalized drug options could be evaluated based on patient-derived organoid cultures obtained from urine (Gijzen et al., 2021; Yousef Yengej et al., 2020) or skin (Barker and Clevers, 2010; Lee and Koehler, 2021) samples with ease. The role of tropomyosin regulation in kidney function (Kobayashi et al., 1983) underscores the importance of the *ex vivo* testing of novel compounds that affect the response to immunotherapy (Augello et al., 2020), such as drugs modulating the activity of GSK3 (Wood, 2022), which regulates cell polarization and migration (Kumar et al., 2009). Agents targeting this enzyme have been implicated in the treatment of a very wide variety of health conditions (Wood, 2022). Our results have implications for the treatment of diseases of the gut as well as the heart. Further investigation using tropomyosin inhibitors could identify the specific mechanisms of cytoskeletal regulation that cells activate in each case when actin function is impaired or altered. Our hypothesis is that targeting actin in resistant tumors could sensitize cancer cells to tubulin inhibitors, which, if proven correct, will have implications in the clinic practice.

The potential clinical application of the analytics approach outlined in this manuscript pertains to anticipating drug resistance (Matov, 2024d) in cancer therapy and the treatment of neurodegeneration. As cell therapy (Gage, 1998) is becoming the forefront of precision medicine, it would be critical to be able to anticipate the mechanisms of action of the engineered (Irvine et al., 2022) immune cells (Matov, 2024e). Natural killer cells and cytotoxic T cells employ different mechanisms to kill their targets (Mitchison, 2021; Topalian and Pardoll, 2025), for instance by secreting cytotoxic lysosomes using the MT cytoskeleton (Blas-Rus et al., 2017) for trafficking and release. There is also a plethora of effects induced in the target cells, most well studied of which lead to apoptosis, necroptosis (Lu et al., 2014), and ferroptosis - a recently identified, potent, caspase-independent mechanism of programmed cell death (Dixon et al., 2012). The exact pathways activated during therapy can be pinpointed by measuring its effects on the target proteins in patient-derived living cells *ex vivo* (Matov, 2024g).

It is conceivable that for different patients, with overall very similar genetic profiles, a distinct type of engineered immune cells would be required. Given the availability of high-quality fundamental research equipment for high resolution live-cell microscopy in most university hospitals, what we propose is to embed in these systems real-time software (Matov, 2024i) for automated image quantification (Matov, 2024h) and on-the-fly statistical analysis of intracellular behavior. Introducing our quantitative imaging method to the clinic would allow physicians to fine-tune, with a high level of certainty, an optimal treatment regimen for each patient.

We propose the utilization of the described cytoskeletal (and DNA fragmentation (Matov, 2024c)) analytics to generate embeddings (Vaswani et al., 2017) for a generative transformer network in which the tokens are the morphology and texture metrics (Matov and Modiri, 2024) of the organoids. The embeddings can include MT and F-actin dynamics before and after treatment with different drugs. We will retrain a transformer network with a new set of tokens - with shape descriptors rather than words and with growth/killing curves (that is, sigmoid curves) rather than sentences of human speech, that is, we will retrain a large language model (LLM) (Elemento et al., 2025) with organoid morphology data. This will allow training of the network to classify each cell as sensitive or resistant and further identify subclasses with different mechanisms of resistance. Further, after training a generative transformer network with organoid morphology and texture as well as live-cell dynamics datasets (Matov, 2024h), we will be able to model disease progression as well.

Reinforcement learning (RL) techniques, such as Markov Decision Processes (Kakade and Langford, 2002; Schneider and Wagner, 1957), can provide the basis for sample stratification based on policies for sample classification in oncology as well as neurodegeneration. In particular, fluid intelligence (Cattell, 1943) applications of language reasoning models that depend on no prior training data can be useful in identifying and diagnosing new clinical conditions in real time. RL is utilized for post-training of LLMs and inference compute. Reasoning or a thought process consists of a sequence of tokens representing the steps in the reasoning trajectories. A Kullback-Leibrer (KL) divergence (Kullback and Leibler, 1951) constraint can achieve good empirical results on a range of challenging policy learning tasks (Schulman et al., 2015). The addition of a Group Relative Policy Optimization (GRPO) (Shao et al., 2024), a modification of the RL algorithm for Proximal Policy Optimization (Schulman et al., 2017), reduces the requirements for training data. For each new sample, GRPO samples a group of outputs from the old policy and then optimizes the policy model by maximizing the objective function, which is the expectation for the average reward of multiple sampled outputs, produced in response to the same question as the baseline. It utilizes a penalty based on computing the Monte Carlo approximations (Kungurtsev et al., 2023) of the KL divergence between the policy model and the reference model.

Our objective has been to present the community with an integrated, easy to use by all, tool (Matov, 2024h; Matov, 2024i) for resolving the complex cellular organization and it is our goal to have such software system approved for use in the clinical practice. A key novelty is that our computational platform outputs results in real-time, during imaging, and without having to store the data first, which will facilitate and significantly speed up research and clinical efforts by providing instant delivery of data analysis. Augmented reality artificial intelligence software (Chen et al., 2019) can be added to live-cell microscopes, thus providing instant visual feedback with added graphics during sample observation. This functionality will enhance the images by also displaying numbers describing the measured on-the-fly differences (Matov, 2024i) and, in doing so, facilitate the image interpretation by the clinical operator. It will significantly enhance the ability of clinicians to make quick and correct conclusions regarding drug action. Our expectation is that improving on our ability to dissect the precise mechanisms of drug action will improve the efficacy of the treatment regimens. Our hypothesis is that targeting actin in resistant tumors could sensitize cancer cells to tubulin inhibitors. If this proves true, it will have implications in the clinic.

## Abbreviations

AMP-PNP: adenylyl imidodiphosphate
AR: androgen receptor
CAF: cancer-associated fibroblast
COX: prostaglandin-endoperoxide synthase
CTGF: connective tissue growth factor
GPX4: glutathione peroxidase, 4
GRPO: group relative policy optimization
GTP: guanosine triphosphate
DoG: difference of Gaussians
EB1: microtubule end binding protein, 1
EB1ΔC: EB1 construct truncated at amino acid 248 that does not interact with other proteins at the microtubule end
EGFP: enhanced green fluorescent protein (2xEGFP indicates two molecules)
FA: focal adhesion
FOLFOX: folinic acid, fluorouracil, and oxaliplatin
FSM: fluorescent speckle microscopy
F-actin: filamentous actin
GSK3α/β: glycogen synthase kinase 3, alpha/beta isoform
HGF: hepatocyte growth factor
KIF11: kinesin-5 protein (homologs – Eg5 in *Xenopus* and KLP61F in *Drosophila*)
LLM: large language model
MRF: Markov random field
MT: microtubule
ORB: oriented FAST (features from accelerated segment test) and rotated BRIEF (binary robust independent elementary features)
PC: prostate cancer
RL: reinforcement learning
ROCK: Rho-associated protein kinase
SIFT: scale-invariant feature transform
SURF: speeded-up robust features
TGFβ: transforming growth factor, beta isoform
Tm2: tropomyosin-2, α-gene
Tm4: tropomyosin-4, δ-gene

## Ethics declaration

IRB (IRCM-2019-201, IRB DS-NA-001) of the Institute of Regenerative and Cellular Medicine. Ethical approval was given.

Approval of tissue requests #14-04 and #16-05 to the UCSF Cancer Center Tissue Core and the Genitourinary Oncology Program was given.

## Data availability statement

The datasets used and/or analyzed during the current study are available from the corresponding author upon reasonable request. The prostate cancer organoid whole genome sequencing datasets (https://www.ncbi.nlm.nih.gov/bioproject/1255230) are accessible at: https://www.ncbi.nlm.nih.gov/sra/SRX28551163.

## Author contributions

A.M. conceived the study, cultured patient-derived organoids and cancer cell lines, performed high resolution microscopy imaging, designed the analysis algorithms, developed the software, performed the analysis, prepared the figures and videos, and wrote the manuscript.

## ACKNOWLEDGEMENTS

I thank Andrea Bacconi for performing perturbation experiments of the long and short isoform of tropomyosin. I am grateful to Jay Gatlin for his feedback to the initial draft of the manuscript and spindle datasets, and Stephanie Gupton, Dylan Burnette, Aaron Groen, Gaudenz Danuser, and Torsten Wittmann for sharing actin and microtubule datasets, and their feedback regarding the shortcomings of the available speckle analysis methodology. I am also grateful to the Genitourinary Tissue Utilization committee and the Genitourinary and Prostate SPORE Tissue Cores at the UCSF Cancer Center for the approval of my tissue requests #14-04 and #16-05, the Stand Up To Cancer / Prostate Cancer Foundation (SU2C/PCF) West Coast Dream Team (WCDT), the Institute of Regenerative and Cellular Medicine for issuing the Institutional Review Board protocol approval IRCM-2019-201, IRB DS-NA-001 for the observational study “Longitudinal analysis of next-generation sequencing of nucleic acids for early detection of degenerative diseases such as cardiovascular, neoplastic and diseases related to the nervous system”, and James Faber for his feedback regarding the protocol and the process of approval.

## SUPPLEMENTARY MATERIALS

Video 1 – Fluorescently-labeled polymerizing MT tips are visualized as EB1 comets forming at the end of the MTs. The comets are detected with ClusterTrack and displayed as green dots. https://vimeo.com/1074106790/83245b9027

Video 2 – ClusterTrack analysis for Vid. 1 is shown in green color for the stages of pausing/fluxing and in red color for the stages of depolymerization. ClusterTrack validation is shown with manual labels for correct (C) or wrong (W) clustering. Forward (F) MT polymerization / EB1 comet tracks are shown in brown color. Observe that several comets are not tracked during polymerization because the brightness of the signal is low and the comets are not detected. https://vimeo.com/1074107156/7022643856

Video 3 – AMPPNP-treated spindle exhibiting no microtubule flux. https://vimeo.com/1018995000/e4e658260e

Video 4 – Monastrol-treated monopole spindle exhibiting inward microtubule flux. https://vimeo.com/1018995640/5ef29f66b9

Video 5 – Flow tracking of two directional microtubule flux in a receptor triple negative MDA-MB-231 breast cancer cell. The main directions are shown as Gaussian distributions at the upper left corner of the image in color-coding. The number of vectors in the flow is reflected in the height of the kernel and the width represents the standard deviation of the direction of motion in each flow. The red vectors track speckles fluxing from the right spindle pole toward the left half of the mitotic spindle. The green vectors track speckles fluxing from the left spindle pole toward the right half of the mitotic spindle. The color of the vectors changes from green to blue (see the second half of the movie) as the spindle rotates, since the color corresponds to the angle of motion. https://vimeo.com/1068615676/e5997e73c1

Video 6 – Growth cone exhibiting a slight contraction in the neck of the cone of the two entangled actin flows. https://vimeo.com/1018995955/42baccdc3f

Video 7 – Flow tracking of four directional actin flux in a PtK1 cell with overexpression of tropomyosin 2. A three-panel movie: The left panel displays the whole cell, the middle panel displays a zoomed-in area in which the vectors from the tracking are showed, the right panel displays a dynamic overlay of the original images with the four overlapped flow fields. Yellow arrows show the retrograde flow field vectors, green arrows show the anterograde flow field vectors, black arrows show the vectors of the upward flow field vectors, and red arrows show the downward flow field vectors. https://vimeo.com/1018996134/0d181b2b08

Video 8 – Tm2 anterograde flow #1. Flow tracking results for the cell shown in Vid. 7 for the major anterograde flow. https://vimeo.com/1068590064/850a021fb3

Video 9 – Tm2 anterograde flow #2. Flow tracking results for the cell shown in Vid. 7 for the minor anterograde flow. https://vimeo.com/1068590379/5ce2315942

Video 10 – Tm2 retrograde flow #1. Flow tracking results for the cell shown in Vid. 7 for the major retrograde flow. https://vimeo.com/1068590778/86e216d026

Video 11 – Tm2 retrograde flow #2. Flow tracking results for the cell shown in Vid. 7 for the minor retrograde flow. https://vimeo.com/1068591097/edef508cc7

Video 12 – Three-directional actin flux in a PtK1 cell with overexpression of tropomyosin 4. https://vimeo.com/1068589045/df2651469c

Video 13 – Cancer-associated fibroblasts in organoid culture 4 days after radical prostatectomy. Patient cells were transduced with an EB1 plasmid on the morning after radical prostatectomy, when the tissue dissociation procedure was over, to fluorescently label polymerizing MT ends. The video was collected during antibiotic selection of fluorescently labeled cells. https://vimeo.com/1043221363/647fea5020

**Figure S1.**
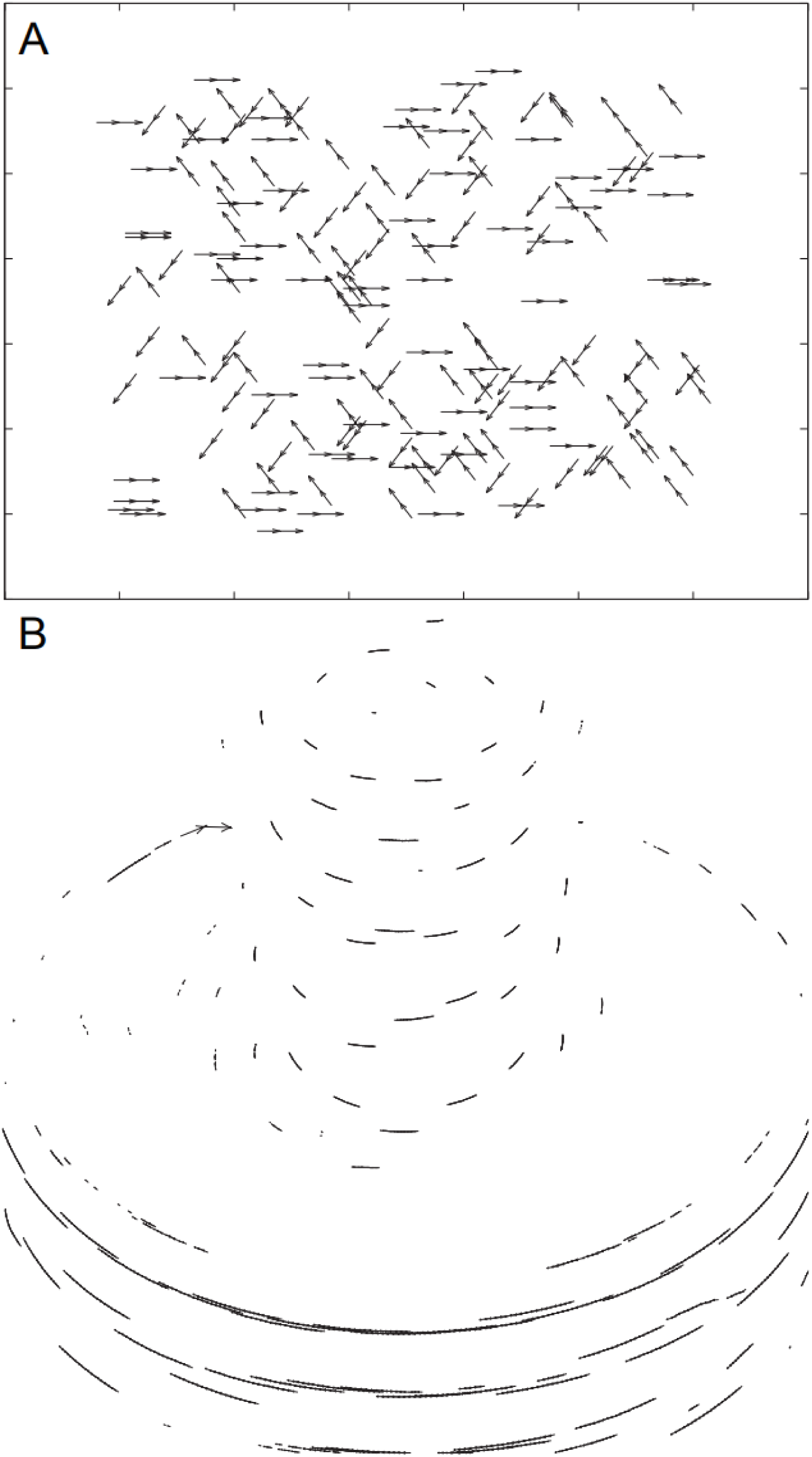
Tracking in artificial datasets representing three-directional motion and circular motion. (A) Tracking with IFTA of artificial data over three time-frames of straight-moving triplets of points moving in three directions, with one of the directions being to the right, another one being toward the upper-left corner of the image, and the third direction being toward the lower-left corner of the image. The tracking algorithm made no linking errors. (B) Tracking with IFTA of artificial data over 20 frames of circular motion of points moving clockwise, with several smaller circles in the upper part of the image and a few larger circles in the lower part of the image. The tracking algorithm made no linking errors. Note that because the motion in this example from one time-frame to the next is relatively slow, most vector arrowheads are very small.

**Figure S2.**
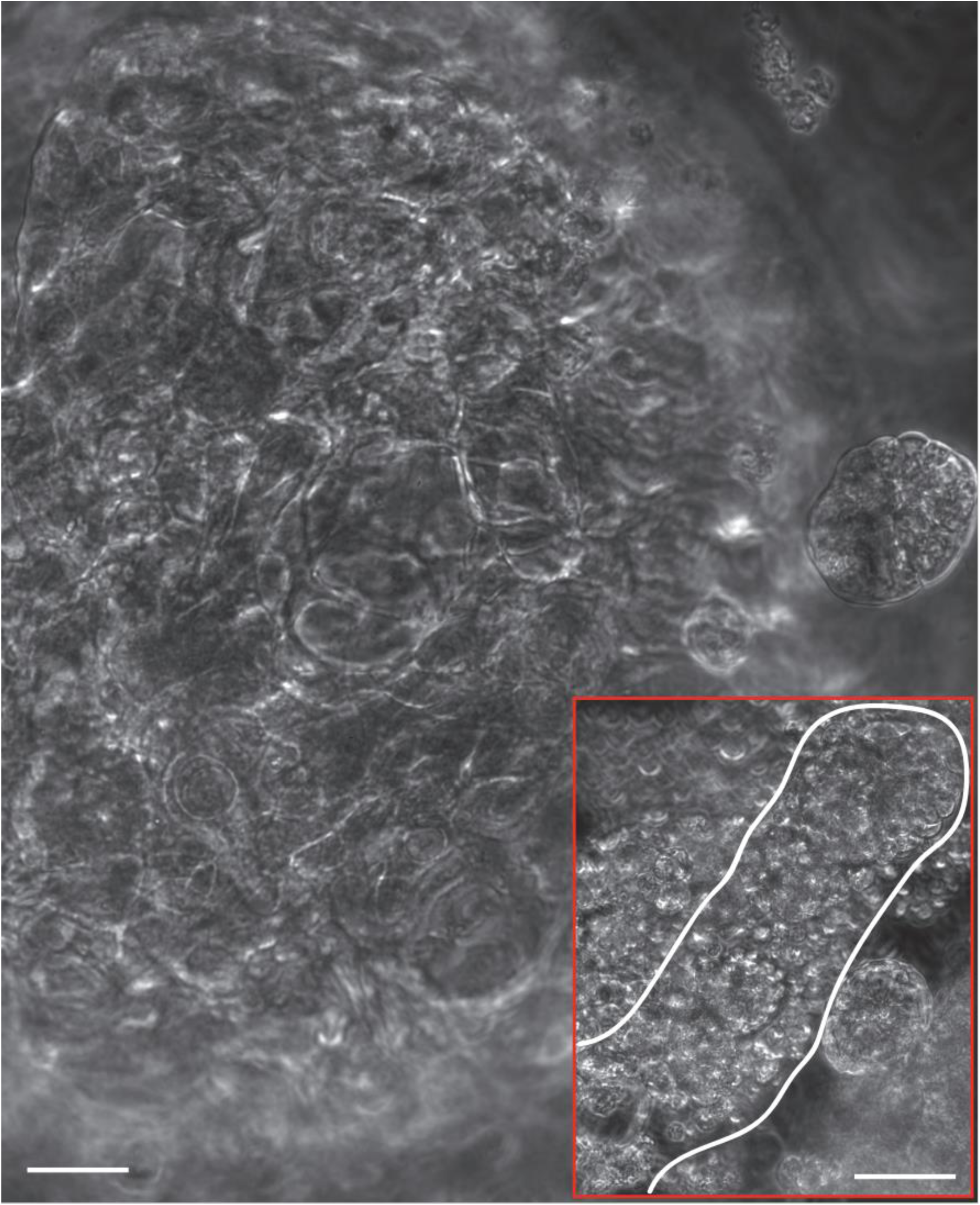
Acinar adenocarcinoma organoids obtained from organ-confined disease. Organoids derived from 1 mg tissue (Gleason Score 5+4, punch core biopsy) from radical prostatectomy. Day 28. The organoids formed visible glandular structures, resembling prostatic ducts (see the inset). Organoid cells formed acini. Phase contrast microscopy, 20x magnification. Scale bar equals 30 µm. Inset scale bar equals 60 µm.

**Figure S3.**
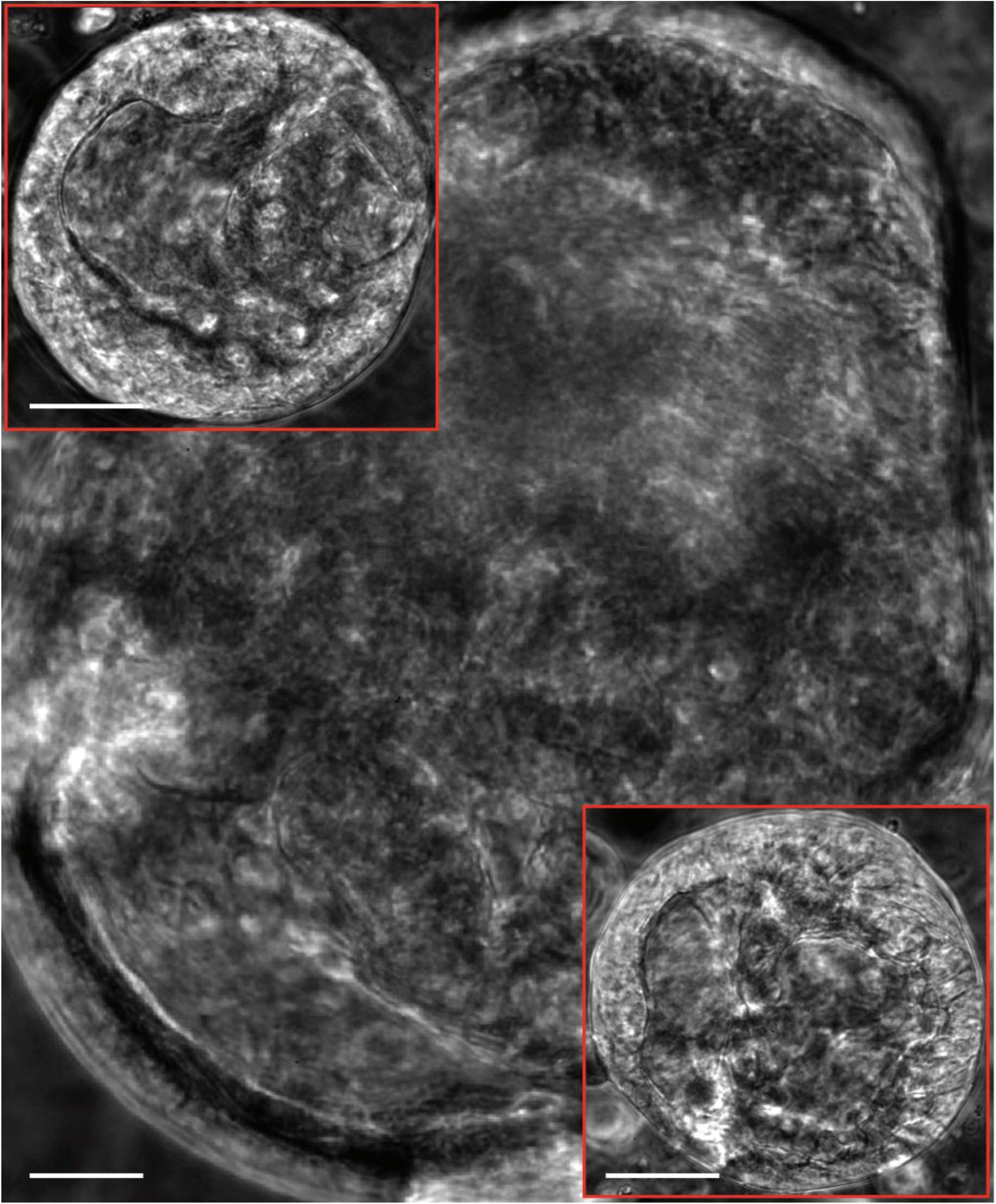
Intraductal prostate cancer organoids obtained from organ-confined disease. Organoids derived from 1 mg tissue (Gleason Score 5+4, punch core biopsy) from radical prostatectomy. Ductal prostate cancer is rare and very aggressive. Day 23. The organoids formed visible ducts and were surrounded by a niche of fibroblasts. Organoids were often shaped like two urethral sinuses. Phase contrast microscopy, 20x magnification. Scale bar equals 30 µm. Insets scale bars equal 50 µm.

**Figure S4.**
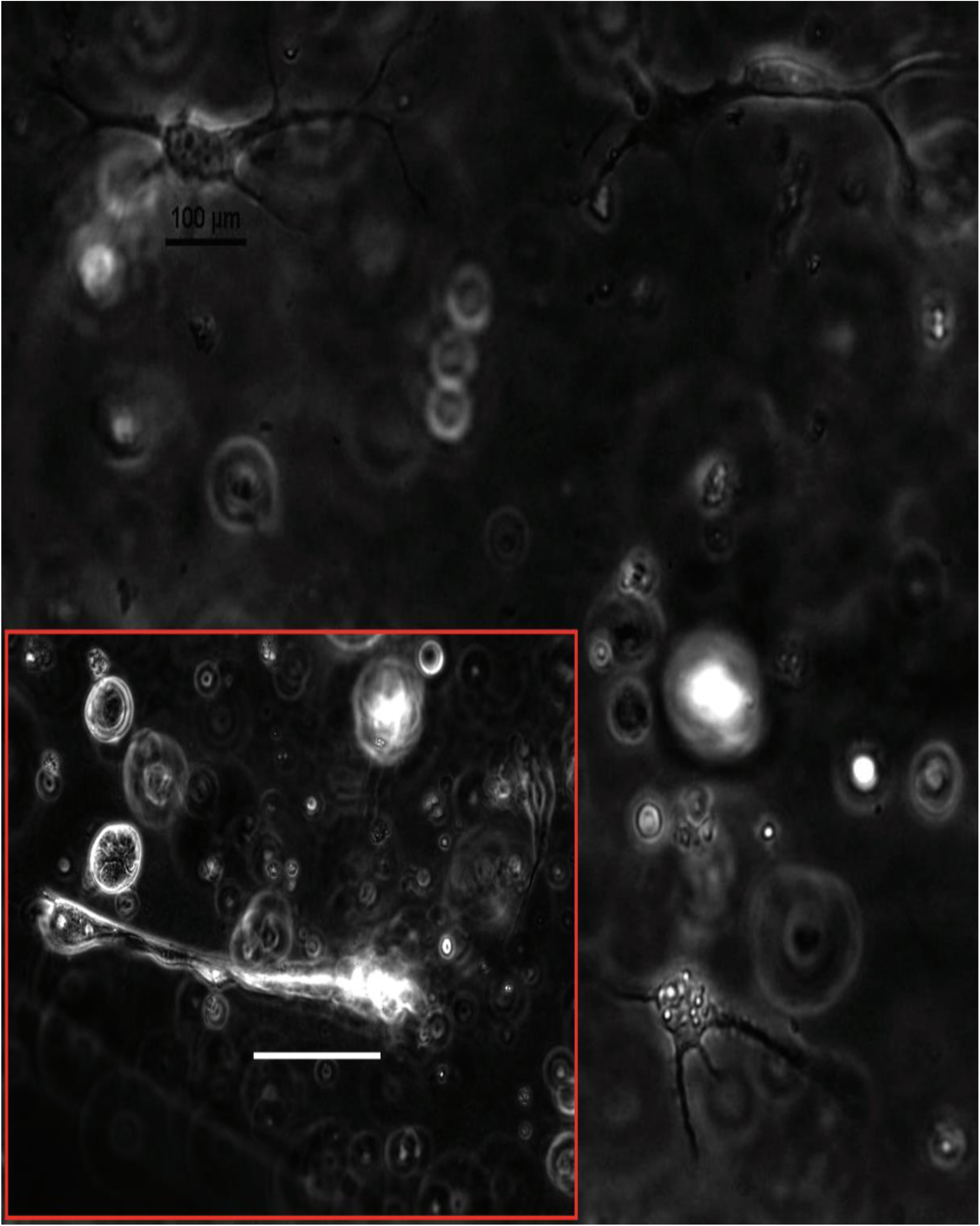
Cancer-associated fibroblasts in samples from radical prostatectomy. Organoids derived from 1 mg tissue (Gleason Score 5+4, punch core biopsy) from radical prostatectomy. Day 2. Organoids begin forming upon fibroblast activation. Inset: Day 4. Phase contrast microscopy, 20x magnification. Scale bars equal 100 µm.

**Figure S5.**
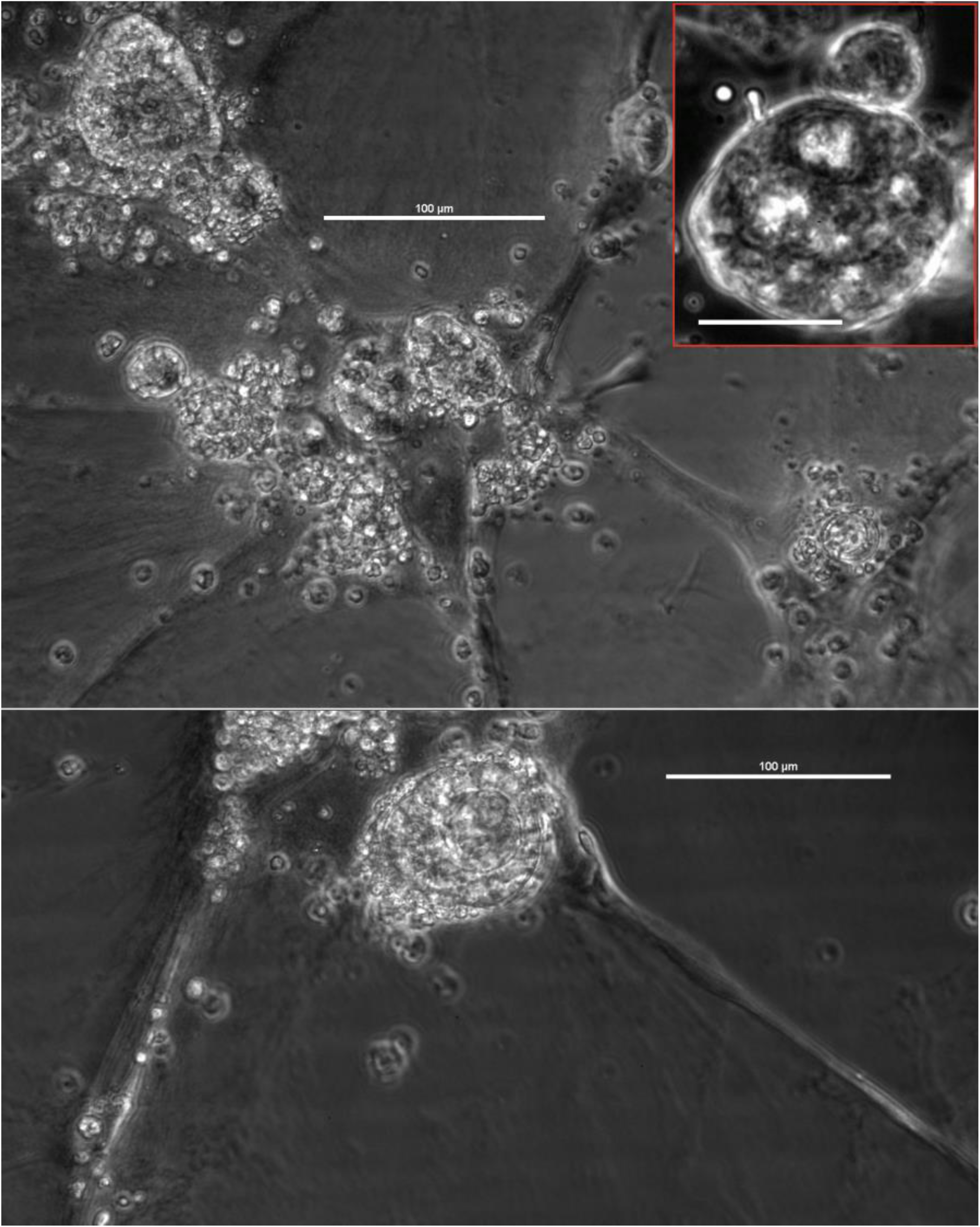
Cancer-associated fibroblasts in samples from retroperitoneal lymph node metastasis. Organoids derived from resected during radical prostatectomy tissue. Drug-naïve tumor. Day 113. Organoids communicate via paracrine signaling. Phase contrast microscopy, 20x magnification. Scale bars equal 100 µm. Inset: Day 2. Scale bar equals 40 µm.

**Figure S6.**
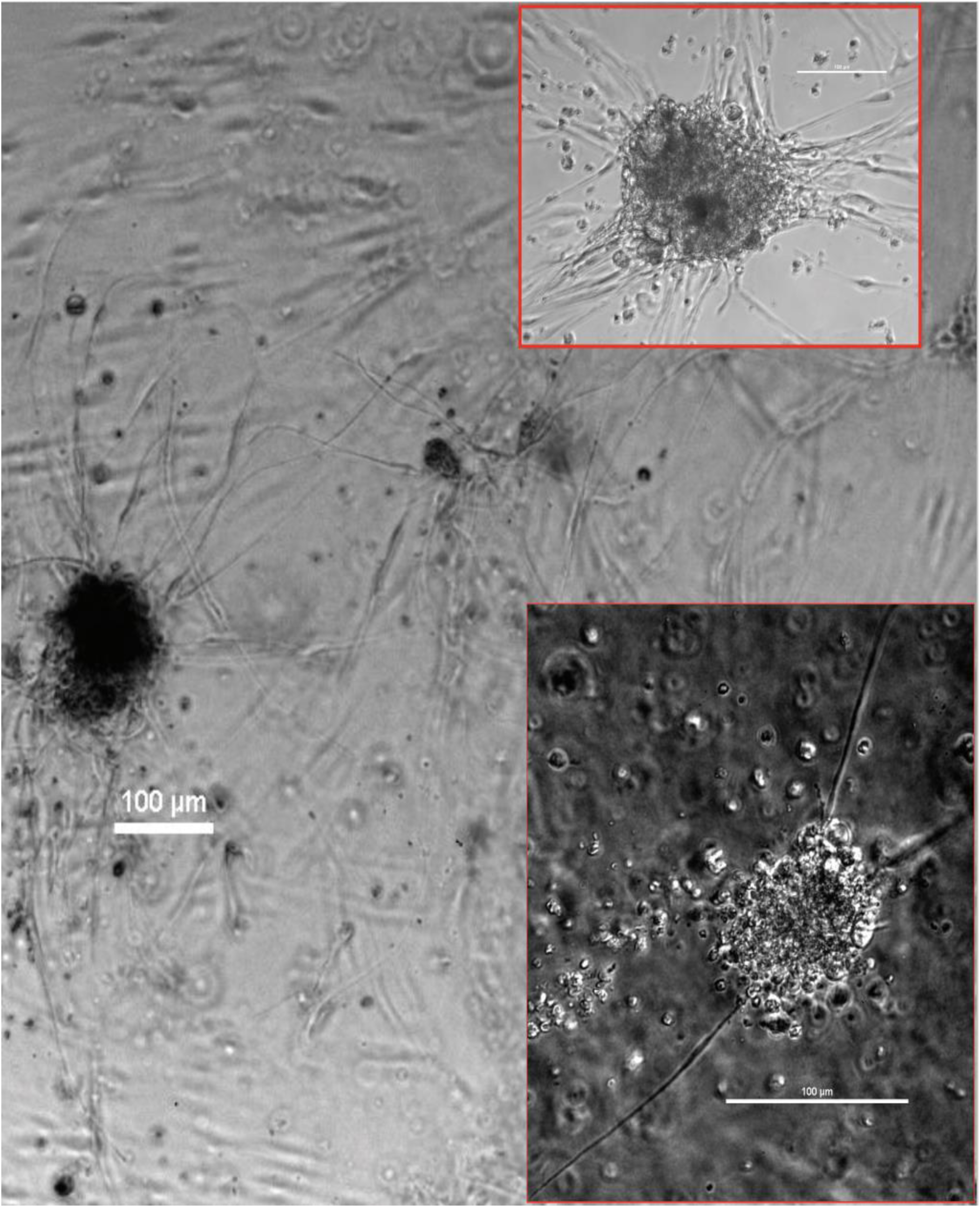
Cancer-associated fibroblasts in samples from retroperitoneal lymph node metastasis. Organoids derived from resected during radical prostatectomy tissue. Prior treatment - androgen deprivation therapy and radiotherapy. Day 84. Transmitted light microscopy, 4x magnification. Upper inset: Day 61. Transmitted light microscopy, 20x magnification. Lower inset: Day 73. Phase contrast microscopy, 20x magnification. Scale bars equal 100 µm.

**Figure S7.**
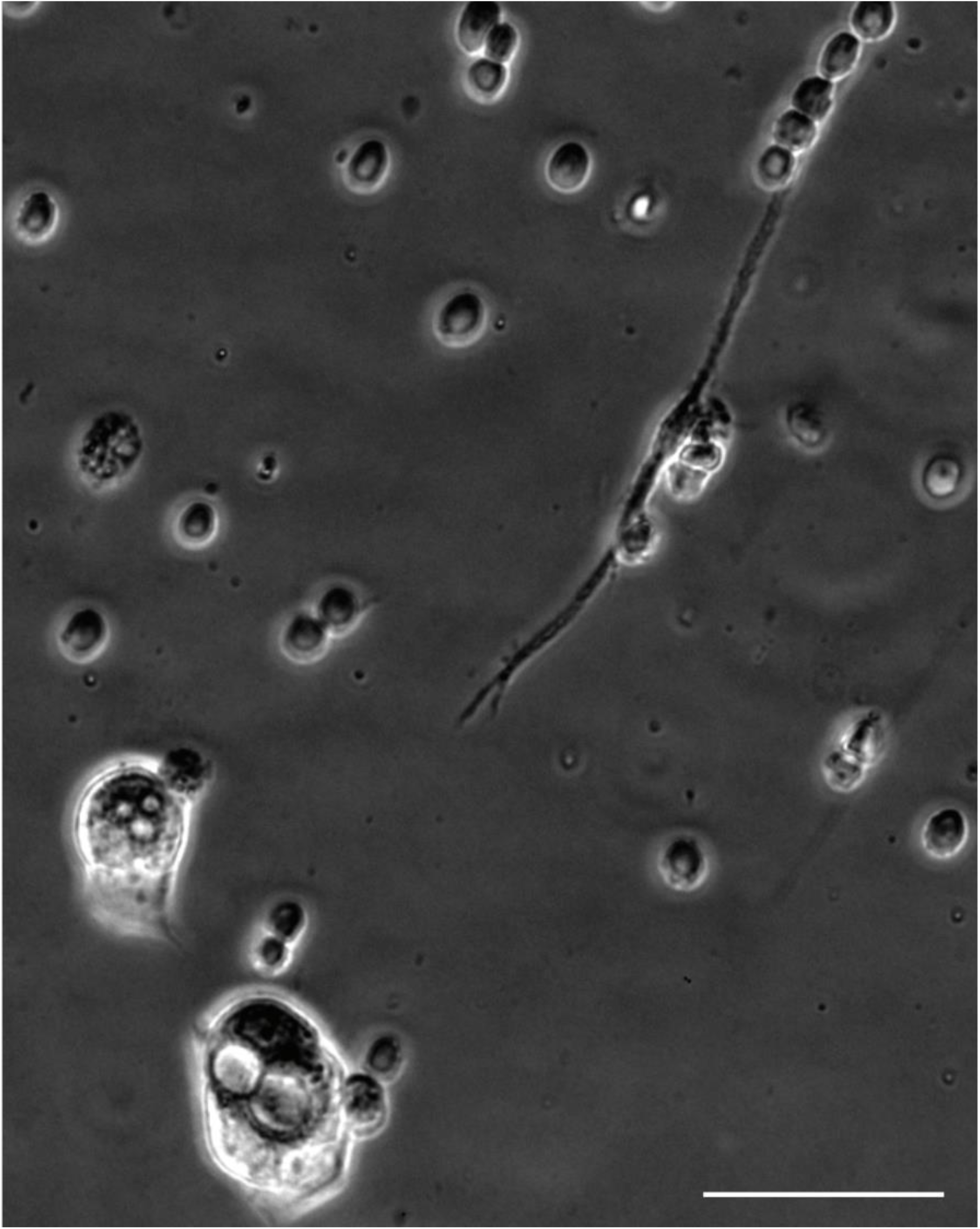
Cancer-associated fibroblasts in samples from needle biopsy of liver metastasis. Organoids derived from ½ mg tissue from needle biopsy. Day 3. Organoids begin forming upon fibroblast activation. Phase contrast microscopy, 20x magnification. Scale bar equals 50 µm.

**Figure S8.**
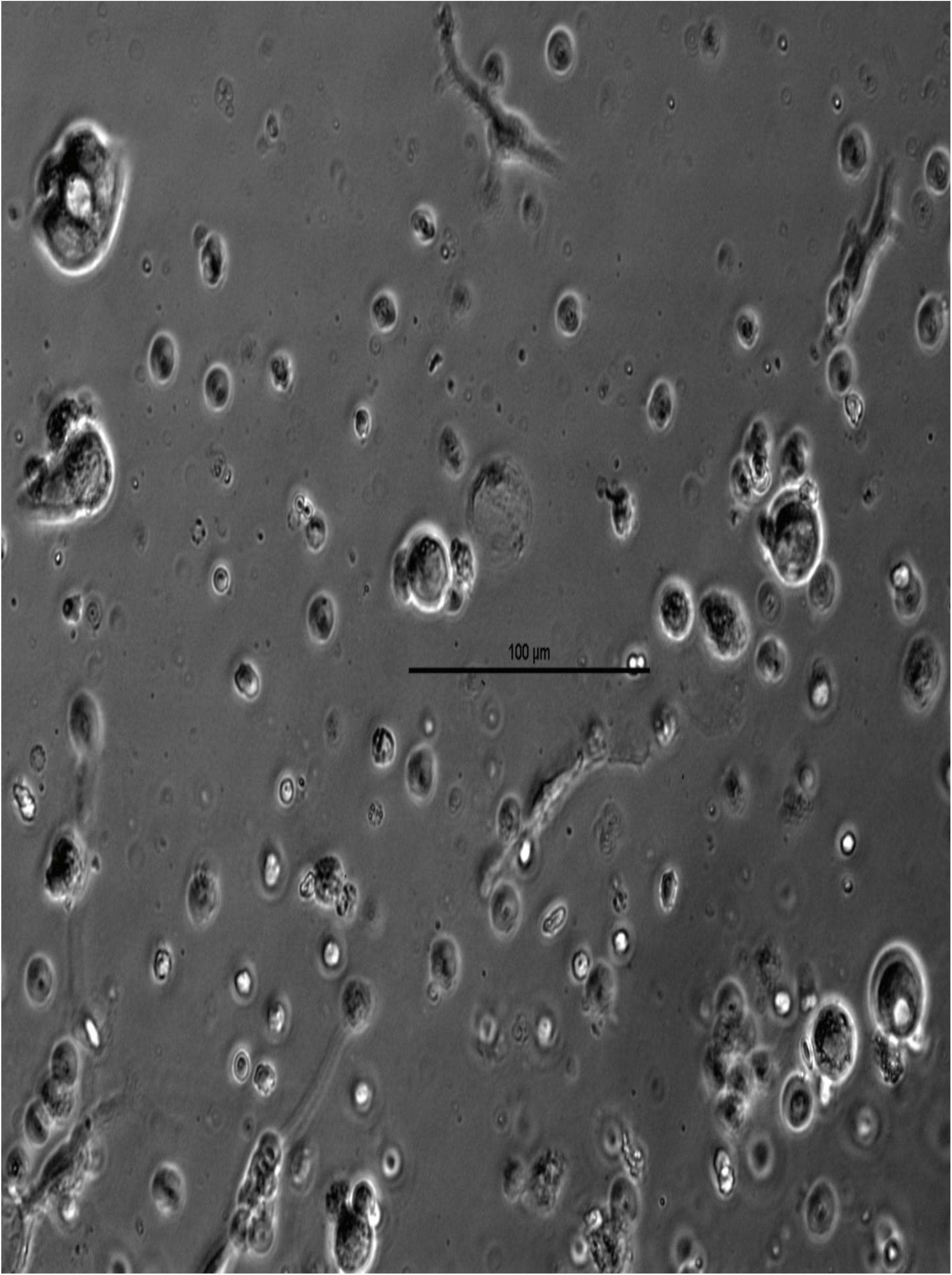
Cancer-associated fibroblasts in samples from needle biopsy of liver metastasis. Organoids derived from ½ mg tissue from needle biopsy. Day 4. Organoids begin forming upon fibroblast activation. Phase contrast microscopy, 20x magnification. Scale bar equals 100 µm.

